# Single-nucleus multiome profiling reveals osteocyte-centered remodeling of human subchondral bone across osteoarthritis severity

**DOI:** 10.64898/2026.10.29.762653

**Authors:** Yutaro Uchida, Hannah Swahn, Yu Okuno, Merissa Olmer, Alia Obeidat, Rachel E. Miller, Anne-Marie Malfait, Miguel Otero, Bella Mehta, Dana E. Orange, William D. Bugbee, The RE-JOIN Consortium, Hiroshi Asahara, Martin K. Lotz

**Author notes:** **Corresponding author:** Martin K. Lotz, MD, Scripps Research Institute, 10550 North Torrey Pines Road, La Jolla, CA 92037.

## Abstract

**Objective:** Subchondral bone remodeling is a hallmark of osteoarthritis (OA), but osteocytes, the principal regulators of bone homeostasis, have been inaccessible to single-cell studies because they are embedded in mineralized matrix. We aimed to define the cellular, transcriptional and chromatin changes of human subchondral bone across OA severity, osteocytes included, and to identify candidate signals between cartilage and bone.

**Design:** Subchondral bone from the medial femoral condyle of human knees graded by the Outerbridge system was analyzed by bulk RNA sequencing (89 knees) and by single-nucleus multiome sequencing of gene expression and chromatin accessibility after mechanical nuclear isolation (16 donors). Three hypotheses were prespecified: with increasing OA severity, (1) the osteoprogenitor fraction increases, (2) the osteocyte fraction increases, and (3) osteocyte SMAD2/3 motif activity increases.

**Results:** With OA severity, bulk expression shifted toward extracellular matrix and ossification programs. The atlas resolved 26 cell states among 61,570 nuclei, including 906 osteocytes. Osteoprogenitors (p = 0.003) and osteocytes (p = 0.036) increased in share along an osteogenic continuum that starts from CXCL12-abundant reticular cells. Chondrocytes at the osteochondral junction upregulated FN1, FGF1, FGF2 and TNFSF11 without increased hypertrophic gene expression, and their share of the inferred FN1 signal received by osteocytes rose from 4% in normal knees to 33% in severe OA. Osteocyte SMAD2/3 motif activity increased with severity (rho = 0.70, p = 0.003), and all three hypotheses remained significant after Holm correction. Across the osteogenic lineage, SMAD2/3 activity correlated with junction chondrocyte FN1 (rho = 0.63, p = 0.017), and in osteocytes it was accompanied by TGF-β-responsive matrix genes and perilacunar-remodeling genes.

**Conclusions:** This osteocyte-inclusive atlas of human OA subchondral bone shows expansion of the osteogenic compartment and a rise of osteocyte SMAD2/3-linked matrix remodeling with disease, and identifies chondrocyte-derived FN1 as a candidate contributor to cartilage–bone communication associated with osteocyte TGF-β/SMAD2/3 signaling.

## Introduction

Osteoarthritis (OA) is now understood as a disease of the whole joint rather than of articular cartilage alone, and subchondral bone is one of its active components contributing to cartilage damage and the generation of joint pain^1,2^. Defining which cells of subchondral bone change, and at which stage of disease, is therefore essential both for understanding the mechanism of OA and for interpreting its significance in disease onset and progression.

The bone changes of OA are neither uniform across the disease course nor simply an increase in bone. Subchondral bone remodeling follows two phases: early disease is dominated by high remodeling with bone loss and thinning of the subchondral plate, whereas thickening and densification of the plate appear only later.^3^ The bone gained in late disease is also of poor quality, since OA cancellous bone is denser than normal bone yet contains proportionally less mineral.^4^ This matters mechanically, because a steep stiffness gradient across the subchondral bone is thought to initiate and advance the overlying cartilage damage.^5^ Understanding OA bone therefore requires measurements that follow disease severity as a graded axis, rather than a single comparison of end-stage “sclerotic” bone with normal bone. The mechanisms proposed for these changes derive largely from mouse models and center on mesenchymal progenitors. After anterior cruciate ligament transection, TGF-β1 is activated in the subchondral bone marrow, inducing clusters of mesenchymal stem cells and marrow osteoid islets, and inhibition of TGF-β activity attenuates degeneration of the articular cartilage.^6^ On the vascular side, type H vessels couple angiogenesis to osteogenesis by accompanying osteoprogenitors,^7^ their excess formation in OA is reduced by TGF-β inhibition,^8^ and preosteoclast-derived PDGF-BB acts as an independent inducer of the same vessel type.^9^ Osteoclast-derived netrin-1, in turn, induces sensory innervation of bone and pain behavior.^10^ In humans, however, these predicted cellular events have not been observed directly: the supporting evidence has so far consisted of tissue TGF-β measurements^6,11^ and histological appearance,^12,13^ not of cell-resolved molecular data.

Human subchondral bone transcriptomics comprises both bulk and single-cell work, but essential constraints remain in each.^14^ Microarray comparison of autopsy control bone with OA bone reported changes in bone-remodeling genes and in WNT and TGF-β/BMP signaling,^15^ and comparison of the degenerated medial with the preserved lateral tibial plateau within the same joint identified expression changes in several hundred genes including *POSTN* and *TNFSF11*.^16^ Paired RNA sequencing of lesional versus preserved subchondral bone from OA knees found genes changing in the same direction in bone and cartilage and proposed *IL11* and *CHADL* as treatment targets,^17^ and pain-associated bone marrow lesions carry a distinct expression signature reflecting osteochondral turnover and inflammation.^18,19^ Because bulk measurements average over the whole tissue, however, they cannot determine which cells carry the changes; and although autopsy and non-OA knees have served as controls,^15,16^ the cohorts have been small and dichotomous, contrasting end-stage arthroplasty tissue with a few autopsy or non-OA surgical knees. How gene expression changes stepwise as grade advances from Normal through early OA to severe OA has therefore remained unknown. Single-cell analysis offers cellular resolution, but the published human series are enzymatic single-cell preparations of arthroplasty tissue, from two to three knee patients^20–22^ or from hips without normal controls,^23,24^ and none recovers the matrix-embedded osteocyte, the most abundant cell in bone. Osteocytes are embedded in mineralized matrix and interconnected through canaliculi, so they are either not released by enzymatic digestion or are destroyed in the process. In the mouse, nuclear isolation of frozen bone recovers osteocytes at an order of magnitude higher yield than enzymatic single-cell preparations,^25^ and single-nucleus chromatin profiling of the marrow stroma resolves fate priming within the osteogenic lineage;^24,26^ neither has been applied to adult human subchondral bone.

We reasoned that single-nucleus multiome sequencing removes these constraints simultaneously: nuclei can be liberated from mineralized tissue by mechanical disruption, frozen specimens can be banked and processed in batch, allowing a cohort that includes donors with macroscopically normal cartilage, and the same nuclei yield both the transcriptome and chromatin accessibility. Here we applied single-nucleus multiome sequencing to human knee subchondral bone from 16 donors spanning macroscopic knee grade 0 to 4, and analyzed in parallel bulk RNA sequencing of 89 donors stratified on the same severity axis. This design resolves osteoblasts, osteoclasts and osteocytes in one dataset and allows cellular composition, cell-cell communication, differentiation trajectories, transcription-factor activity and chromatin accessibility to be interrogated stepwise along the severity axis from Normal to severe OA. We report that the osteogenic lineage progressively activates bone-matrix production and ossification programs and its osteoprogenitors and osteocytes occupy a larger share of the sampled nuclei, that the osteocyte sits at the center of OA signaling as a sender of TGF-β and a receiver of *FN1* whose chondrocyte-derived share rises with grade, with chondrocyte FGF directed at the osteoprogenitor, and that on chromatin SMAD2/3 motif activity rises across the chondro-osteogenic axis and converges, in the osteocyte, on matrix and perilacunar-remodeling target genes.

## Results

### Overview of study design

Entire knees were obtained from organ donors and staged on the Outerbridge macroscopic knee grade,^27^ based on scoring all cartilage surfaces of the knee, and classified as Normal (grade 0-1), early OA (grade 1.5-2) or severe OA (grade 3-4). Osteochondral tissues were resected from the weight-bearing region of the medial femoral condyle and cartilage and bone were separated for the analyses. Osteochondral tissues were also obtained from OA patients undergoing total knee arthroplasty. Subchondral bone was analyzed by bulk RNA-seq of 89 donors (20 Normal, 16 early OA, 53 severe OA) and by single-nucleus multiome sequencing, profiling RNA and ATAC in the same nuclei, of 16 donors (6 Normal, 3 early OA, 7 severe OA). From these data we identified differentially expressed genes (DEGs), performed gene set enrichment analysis (GSEA), inferred cell-cell communication with CellChat, analyzed differentiation of the bone-related cells by pseudotime analysis, and measured transcription-factor motif activity on chromatin (Fig. 1a, Supplementary Tables 1-2). Histopathological changes were assessed with the OARSI scoring for cartilage^28^ and the osteoarthritis bone score (OABS) for subchondral bone;^29^ representative sections across the OARSI and OABS range are shown in Fig. 1b.

**Figure 1.**
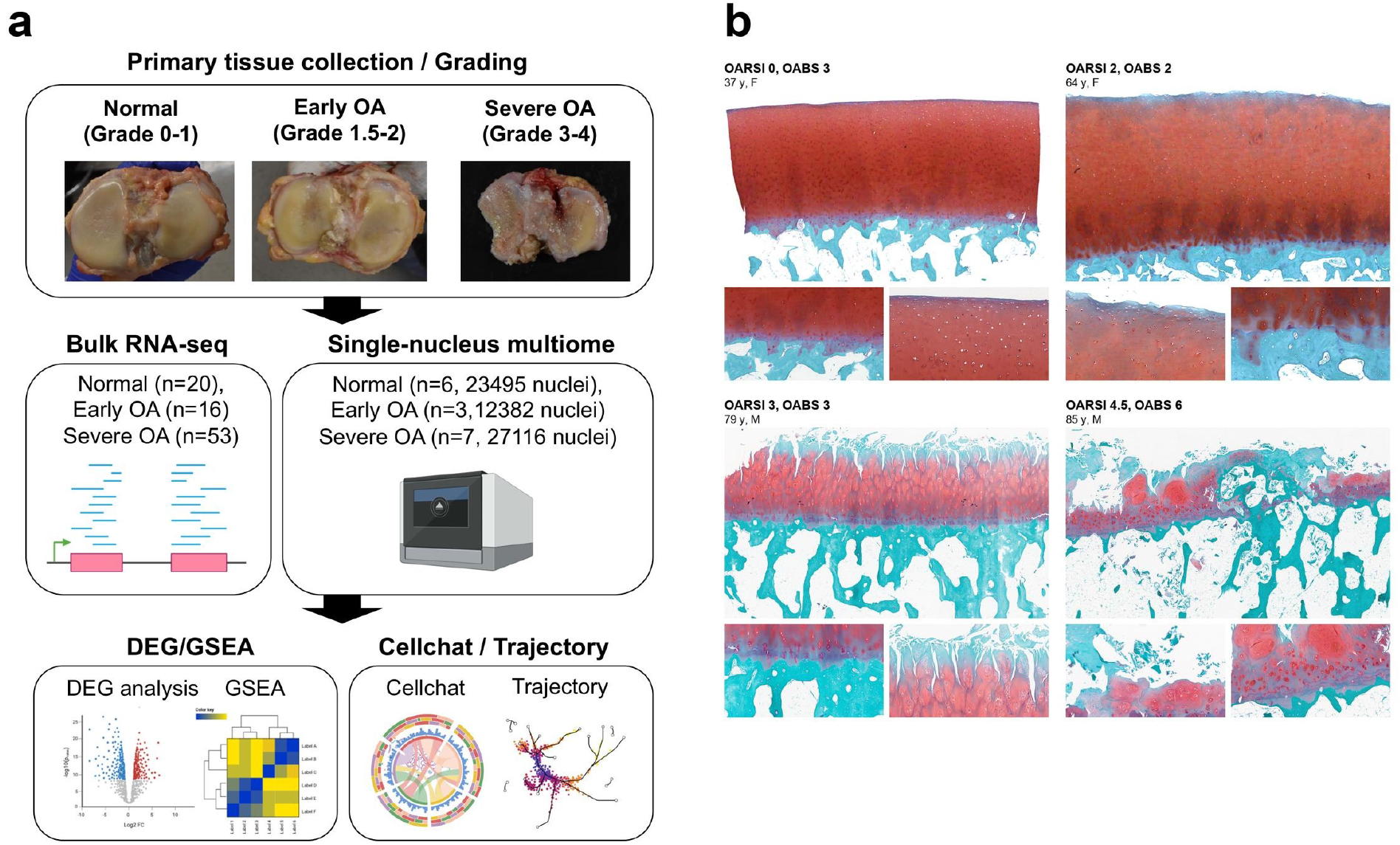
Study design and the structural state of the osteochondral unit. (a) Primary tissue collection and grading: representative macroscopic images of the articular surfaces for Normal (macroscopic knee grade 0 to 1), early OA (grade 1.5 to 2) and severe OA (grade 3 to 4; Outerbridge). The two sequencing arms with their donor numbers, bulk RNA-seq (20 Normal, 16 early OA, 53 severe OA) and single-nucleus multiome sequencing (6, 3 and 7 donors; 23,495, 12,382 and 27,116 nuclei passing the nucleus-level quality filter, of which 61,570 enter the atlas after the chondrocyte transcript filter), and the analysis workflow: differential expression and gene set enrichment, cell-cell communication (CellChat), trajectory analysis and chromatin accessibility. (b) Representative Safranin O and fast green sections of the osteochondral unit from four donors (OARSI score 0, 2, 3 and 4.5; OABS 3, 2, 3 and 6; age and sex as labeled): an overview of cartilage and subchondral bone (top) with a mid-power field of the cartilage and a high-power field of the osteochondral junction (below), illustrating the appearance of the osteochondral unit across OARSI and OABS scores. The molecular analyses of Figures 2 to 8 run on the macroscopic knee grade of the same knees.

### Bulk transcriptome analysis reveals upregulation of ECM and downregulation of gene expression regulatory genes with OA progression

To identify genes whose expression changes linearly with advancing OA in subchondral bone, we tested bulk RNA-seq data of subchondral bone from the 89 knees against continuous macroscopic knee grade. Principal component analysis of these donors showed that Normal and severe OA samples occupy different regions of the first two components (Supplementary **Fig. 1**; PC2, Wilcoxon *p* = 5.0e-07). Of 17,366 genes tested, 491 changed significantly, 224 rising and 267 falling with grade (**Fig. 2a**, Supplementary Table 3). Gene set enrichment placed extracellular matrix (ECM) organization, collagen fibril organization, collagen metabolic process, skeletal system development and biomineral tissue development among the positively regulated Gene Ontology terms, while the negatively regulated terms comprised cytoplasmic translation, ribonucleoprotein complex biogenesis, ribosome biogenesis, RNA splicing and nucleosome organization (**Fig. 2b**). Among the leading-edge genes, the ECM components *COL3A1*, *COL5A1* and *COL6A2* rose linearly with advancing grade, whereas *EIF3H*, *NPM1* and *HSP90AB1*, regulators of gene expression, declined (Fig. 2c, Supplementary Table 4). The significant genes also included components of the TGF-β and FGF axes: *TGFB1*, *LTBP2*, *LTBP3* and *FGFR1* rose, *TGFBR3* fell, and the TGF-β-responsive matrix genes *COL1A1*, *COL1A2*, *BGN* and *FN1* rose with *COL3A1* (**Fig. 2a**).

**Figure 2.**
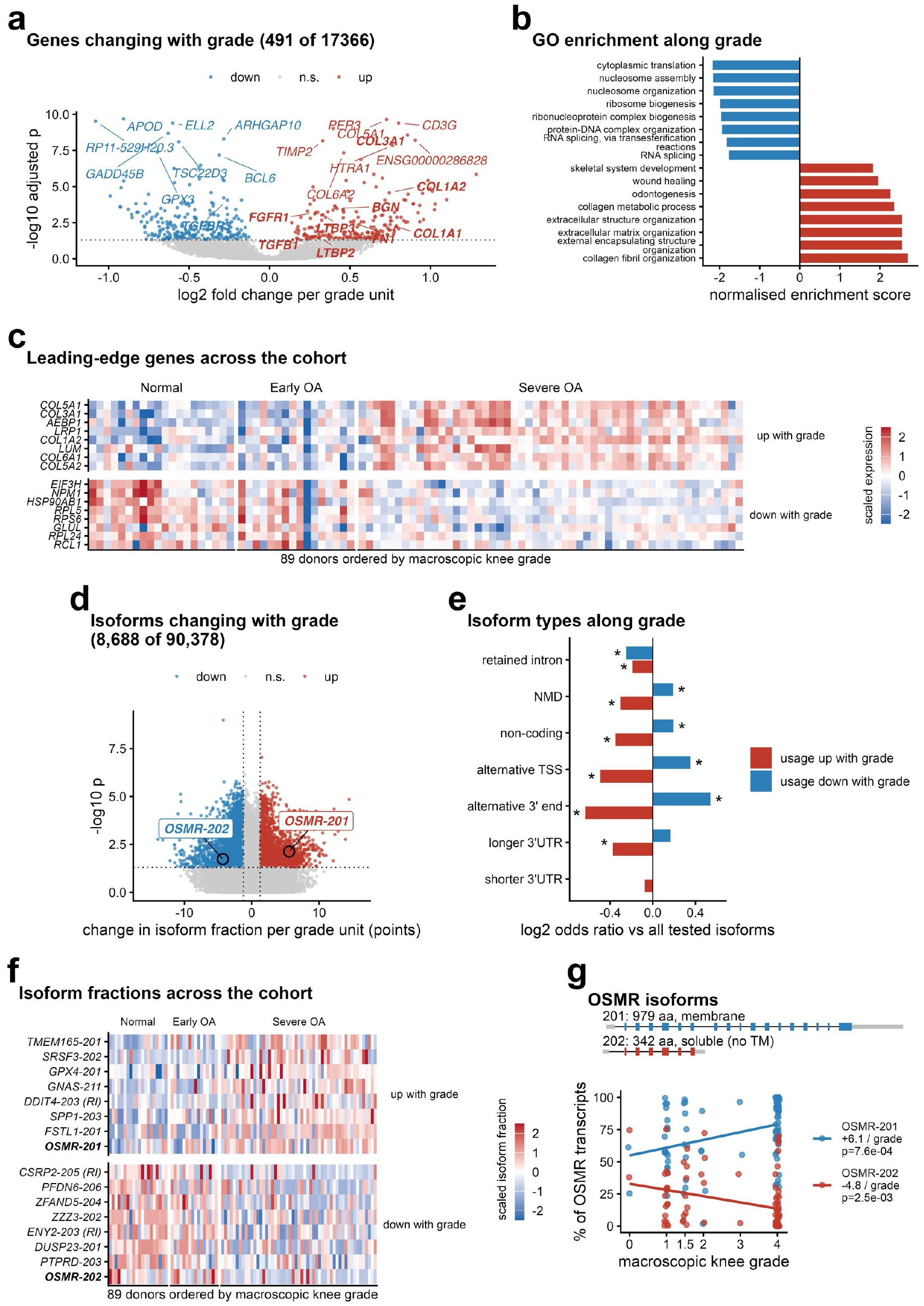
The tissue-level response to severity in bulk RNA-seq: gene expression and isoform usage. (a) Genes changing linearly with grade in bulk RNA sequencing of 89 donors; DESeq2 on continuous grade with batch and a detected-genes covariate. The eight genes with the smallest adjusted p values in each direction are labeled, red for rising and blue for falling genes; the significant genes of the TGF-β and FGF axes (pre-specified list in Methods) are labeled in bold. (b) Gene set enrichment on the same ranking (GO biological process), the eight strongest terms in each direction; the full significant list is in Supplementary Table 4. (c) Protein-coding leading-edge genes of the strongest program in each direction, as scaled expression with one column per donor, the 89 donors ordered by macroscopic knee grade and grouped by stage. (d) Isoforms whose usage changes with grade in the same 89 knees: the change in isoform fraction per grade unit (percentage points; linear model with batch and the detected-genes covariate) against the p value of the partial Spearman correlation of the fraction with grade; 8,688 of 90,378 isoforms at nominal p < 0.05 and at least 1.25 points per grade unit, 4,311 of them also at adjusted p < 0.05, red rising and blue falling. The two *OSMR* isoforms are marked. (e) Enrichment of isoform classes among the rising and the falling isoforms relative to all tested isoforms (log2 odds ratio, Fisher’s exact test; asterisk p < 0.05); classes are defined against each gene’s principal isoform in normal knees (Methods). (f) Isoform fractions of the seven strongest rising and falling isoforms whose usage is spread across the cohort, with the two *OSMR* isoforms, scaled per isoform, one column per donor ordered by macroscopic knee grade and grouped by stage. (g) *OSMR*: exon structure of the membrane (201) and soluble (202) isoforms, and the share of *OSMR* transcripts carried by each per knee against grade, with linear fits. Details in Methods.

The same 89 knees also show changes in which isoform of a gene is used. From the alignments we quantified 90,378 isoforms of 14,100 genes against the GENCODE v44 annotation and tested the fraction of each gene’s output carried by each isoform against grade, with the covariates of the gene-level model (Methods). The usage of 8,688 isoforms in 6,029 genes changed with grade, 3,412 rising and 5,276 falling by at least 1.25 percentage points per grade unit, 4,311 of them also below the Benjamini-Hochberg threshold (Fig. 2d, Supplementary Table 5). Relative to all tested isoforms, the falling isoforms were enriched for transcripts with an alternative transcription start or 3′ end, for non-coding transcripts and for transcripts subject to nonsense-mediated decay, and the rising isoforms were depleted of the same classes (Fig. 2e); with advancing OA, usage converges on each gene’s principal transcript. Among the isoforms whose usage shifts across the whole cohort (Fig. 2f) are the two transcripts of the oncostatin M receptor gene *OSMR*: the transcript encoding the membrane receptor (OSMR-201) gained 6.1 points of *OSMR* usage per grade unit, and the transcript that ends within intron 7 and encodes the soluble receptor without a transmembrane segment (OSMR-202) lost 4.8 points (Fig. 2g), with total *OSMR* expression unchanged (log2 fold change 0.02 per grade unit; Supplementary Table 3).

### The tissue response tracks a bone histology score as well as the macroscopic grade

The macroscopic knee grade scores the cartilage surfaces of the whole joint, whereas the bone was taken from one region of the medial femoral condyle. We therefore repeated the bulk analysis against the OABS of the subchondral bone of that region, available for 85 of the 89 knees, under the model used for the grade. One hundred and twenty-seven genes changed with the OABS, 56 rising and 71 falling (Supplementary Fig. 2, Supplementary Table 3). Of the 491 genes that change with grade, 471 moved in the same direction with the OABS, 280 reached nominal significance and 55 the adjusted threshold, and the two sets of fold changes correlated at r = 0.85; across all 17,366 tested genes the two axes correlated at r = 0.61. Within the 48 arthroplasty knees, where the macroscopic grade is uniform and the OABS score ranges from 1 to 7, the severity signature of the 491 grade genes rose with the OABS score (rho = 0.42, p = 0.003; rho = 0.39 adjusting for age, sex, BMI, post-mortem interval, batch and detected genes).

### Single-nucleus analysis of subchondral bone identifies 26 cell states including osteocytes and osteoclasts

To capture the gene expression changes of OA progression at single-cell resolution, we profiled the subchondral bone of the 16 donors and analyzed 61,570 nuclei (Methods). Clustering resolved 26 cell states falling into five lineage groups, annotated by canonical marker genes: myeloid/lymphoid cells, vascular/perivascular cells, marrow stroma, a chondrogenic state, and the osteogenic lineage (**Fig. 3a**); nuclei counts and marker expression per state are shown in **Fig. 3b** and **3c**. The myeloid and lymphoid group comprises resident macrophages (10,250 nuclei; *LYVE1*, *MRC1*, *CD163*), inflammatory macrophages (1,759; *IL1B*, *NLRP3*), monocytes (1,150; *FCN1*, *VCAN*), dendritic cells (1,368; *FLT3*, *HLA-DQA1*), cycling myeloid cells (1,035; *MKI67*, *TOP2A*), osteoclasts (445; *CTSK*, *ACP5*), T and NK cells (3,250; *CD3E*, *NKG7*), B cells (397; *MS4A1*, *CD79A*) and plasma cells (500; *MZB1*, *JCHAIN*). The vascular and perivascular group comprises arterial endothelial cells (ECs) (1,342; *GJA5*, *SEMA3G*), capillary ECs (1,292; *RGCC*, *CA4*), sinusoidal ECs (1,629; *STAB1*, *STAB2*), Type-H ECs (297; *EMCN*), angiogenic ECs (1,798; *APLN*, *ESM1*), lymphatic ECs (1,810; *PROX1*, *PDPN*), and pericytes/mural cells (1,734; *ACTA2*, *NOTCH3*). The marrow stroma comprises the *CXCL12*- and *LEPR*-expressing CAR cell (4,664),^30^ activated fibroblasts (1,960; *TNFAIP6*, *COL3A1*), stress fibroblasts (740; *NR4A1*, *FOSB*) and adipocytes (8,472; *GPAM*, *PLIN1*). The chondrogenic state is represented by a single population, the junction chondrocyte, the chondrocytes of the calcified and deep cartilage retained with the bone cores (3,979 nuclei after the exclusion of 1,423 nuclei that carried erythroid, plasma-cell or myeloid programs or almost no chondrocyte transcripts; *ACAN*, COL2A1; Methods, Supplementary Table 14). The osteogenic lineage is resolved into five states: the osteoprogenitor (6,354; *WIF1*, *CDH11*), the *WIF1*+ osteoblast (2,328; *RUNX2*, *SP7*), the stromal osteoblast (757; *COL6A3*, *COL16A1*), the mature osteoblast (1,354; *COL1A1*, *IBSP*, *PRSS35*) and the osteocyte (906; *DMP1*, *MEPE*, *CD44*, *SOST*, *PHEX*). This analysis therefore captured the cell populations that are difficult to recover by single-cell approaches, including the osteocyte and the osteoclast.

**Figure 3.**
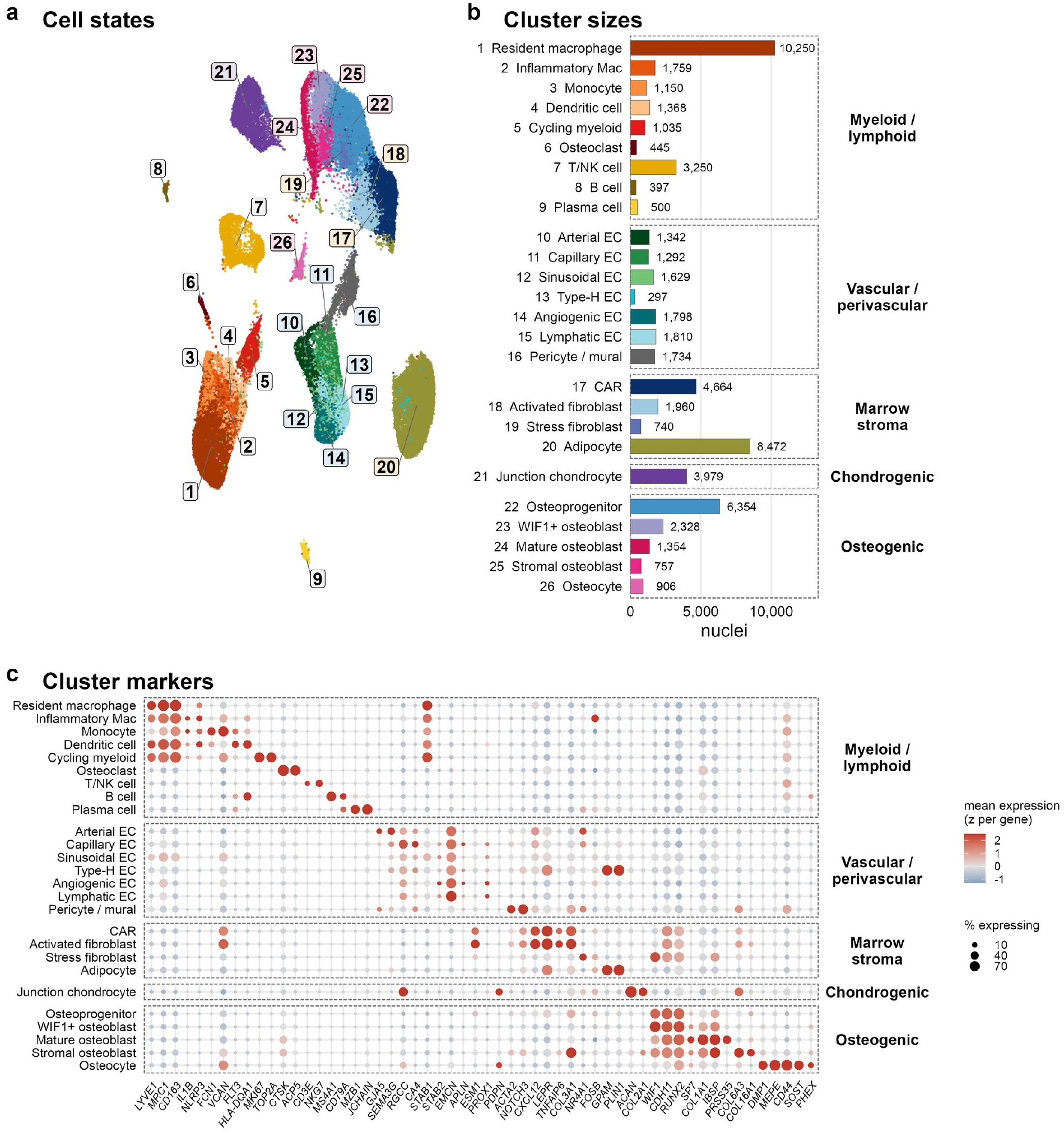
Single-nucleus atlas of human subchondral bone. (a) UMAP of 61,570 nuclei from 16 donors, colored by cell state. (b) Nuclei per state, grouped by lineage; the junction chondrocyte count is after the exclusion of 1,423 non-chondrocyte nuclei (Methods). (c) Marker expression per state: canonical anchors plus literature-audited data-driven markers, lineage-ordered. Details in Methods.

### Bone matrix production and ossification increase in the osteogenic clusters and the junction chondrocyte with OA progression

We next analyzed, for each cell type, the genes changing linearly with OA progression, testing each cluster’s pseudobulk profile against continuous grade with a capture-burden covariate. Across the atlas, 5,654 significant gene-by-cluster associations were found (each gene counted once per cluster in which it changed, across the 25 cell states that clear the donor threshold) (**Fig. 4a**, Supplementary Figs. 3-6, Supplementary Table 6). Gene set enrichment analysis (GSEA) of these expression changes showed that ECM organization and glycoprotein biosynthesis rose in the stromal osteoblast and cartilage development in the osteoprogenitor (Fig. 4b-e), and endochondral ossification in the WIF1+ osteoblast, whose enrichment is given in Supplementary Table 7. At the level of individual genes, in osteocytes 10 genes rose with OA progression, including TNFRSF11B, known as osteoprotegerin (OPG), together with SMOC2 and ECRG4, while 6 genes fell, including TLR4, SLC1A1 and LAMA2 (Fig. 4f). In the osteoprogenitor, 79 genes rose, including *ECRG4*, *EPAS1* and *TNFRSF11B*, and 137 fell, including PDK3, CTSB and PTX3 (Fig. 4g). In the WIF1+ osteoblast, 24 genes rose, including *CILP*, *SETBP1* and *FMOD*, and 66 fell, including PTX3, FYN and PTPN11 (Supplementary Fig. 6). In the stromal osteoblast, 310 genes rose, including OGN, COLGALT2 and COL2A1, and 234 fell, including TWIST1, DDIT4 and PTX3 (Fig. 4h). In the junction chondrocyte, 211 genes rose and 264 fell: the rising genes included FN1 (log2 fold change +1.01 per grade unit), FGF1 (+1.00), FGF2 (+0.59), TNFSF11 (+1.10), COMP (+0.86), HTRA1 (+0.99), CLEC3A (+1.21), LUM (+0.98), ECRG4 (+0.88), FMOD (+0.76), INHBA (+0.94) and TNFRSF11B (+0.67), the falling genes *ADGRD1*, *MAOA*, *ARID3A*, *DDIT4*, *ACSL1* and *ABCA1*, and on its grade ranking cartilage development (NES 2.28), skeletal system development (1.99), ECM (1.78) and ossification (1.64) rose while fatty acid metabolism fell (Fig. 4e, i, Supplementary Tables 6 and 14).

**Figure 4.**
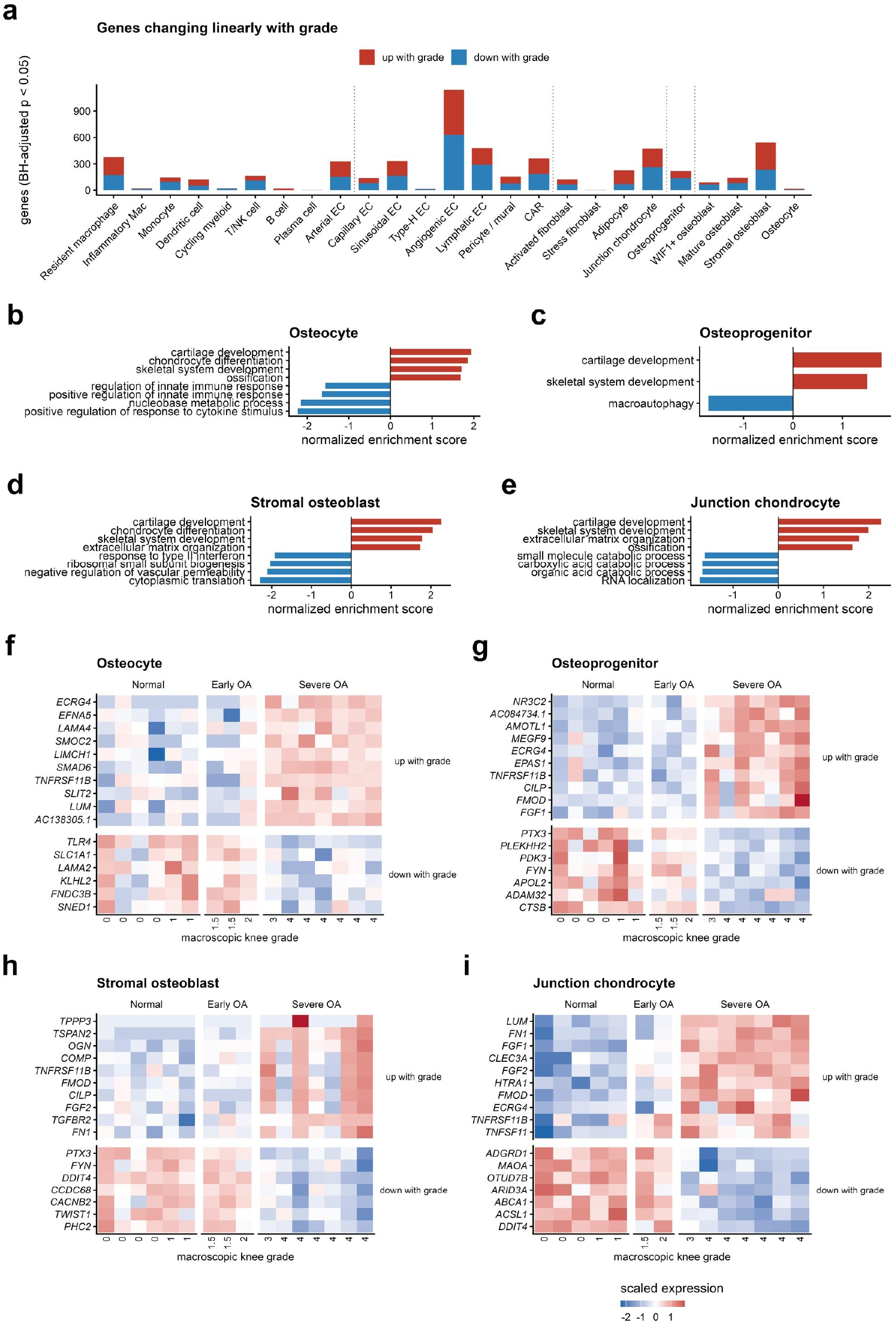
Genes changing linearly with grade, per cell state. (a) Number of genes changing linearly with macroscopic knee grade within each of the 25 cell states that clear the donor threshold, from the per-state pseudobulk model with the capture-burden covariate; dotted lines separate the lineage groups. (b to e) GSEA (GO biological process) on the continuous-grade ranking of the osteocyte, the osteoprogenitor, the stromal osteoblast and the junction chondrocyte; bars are normalized enrichment scores of significant terms (BH-adjusted *p* < 0.05), up to four per direction with skeletal-program terms shown first. The full significant lists are in Supplementary Table 7. (f to i) Rising and falling genes of the same four states, shown as that state’s own pseudobulk: the characterizing genes of each state and the significant genes of the TGF-β and FGF axes (*CILP*, *FMOD*, *COMP*, *FN1*, *FGF1*, *FGF2*, *TGFBR2*, *TNFRSF11B*), together with the genes with the smallest adjusted p values. Each column is a donor, labeled with that donor’s grade and grouped by stage; expression is scaled per gene. The WIF1+ osteoblast, the mature osteoblast and the states outside the osteogenic lineage are in Supplementary Figures 3 to 6. Details in Methods.

### Chromatin accessibility reveals a coordinated gain of SMAD2/3 motif activity across the chondro-osteogenic axis

We next measured transcription-factor family motif activity in the cell states of the same nuclei by ATAC-seq, comparing severe OA against Normal and early OA donors (nominal Wilcoxon p; the BH-adjusted values and the Spearman correlation with continuous grade are given alongside in Supplementary Table 12). Across the tissue, the adipogenic CEBP family fell in nearly every compartment, immune and vascular states included, whereas SMAD2/3 motif activity rose as a feature shared by the osteogenic clusters, with nominally significant gains across the chondro-osteogenic axis (junction chondrocyte +1.16, p = 0.004; CAR +0.88, p = 0.005; osteoprogenitor +0.79, p = 0.042; WIF1+ osteoblast +1.00, p < 0.001; stromal osteoblast +1.11, p = 0.002; osteocyte +0.79, p = 0.003) (**Fig. 5a**, Supplementary Table 12). The BMP-arm SMAD1/5 rose in the stromal osteoblast (+0.91, p = 0.042), the adipocyte (+1.08, *p* = 0.012) and the resident macrophage (+1.07, *p* = 0.008). Stage-resolved heatmaps of the junction chondrocyte and the osteogenic clusters confirm the progressive gain of SMAD2/3 activity as OA advances (Fig. 5b). In the junction chondrocyte the NFAT family rose alongside SMAD2/3 (+1.17, p = 0.002) while the EGR, CEBP and TEAD families fell (Fig. 5b), and at the level of individual peaks the 29,870 peaks that opened with grade were enriched for NFAT (1.76-fold) and SMAD2/3 (1.29-fold) motifs and the 1,985 peaks that closed for KLF/SP, EGR and CEBP motifs (Supplementary Table 14). The chromatin of the osteogenic clusters in advancing OA is thus characterized by a loss of the CEBP program and a gain of SMAD2/3 activity.

**Figure 5.**
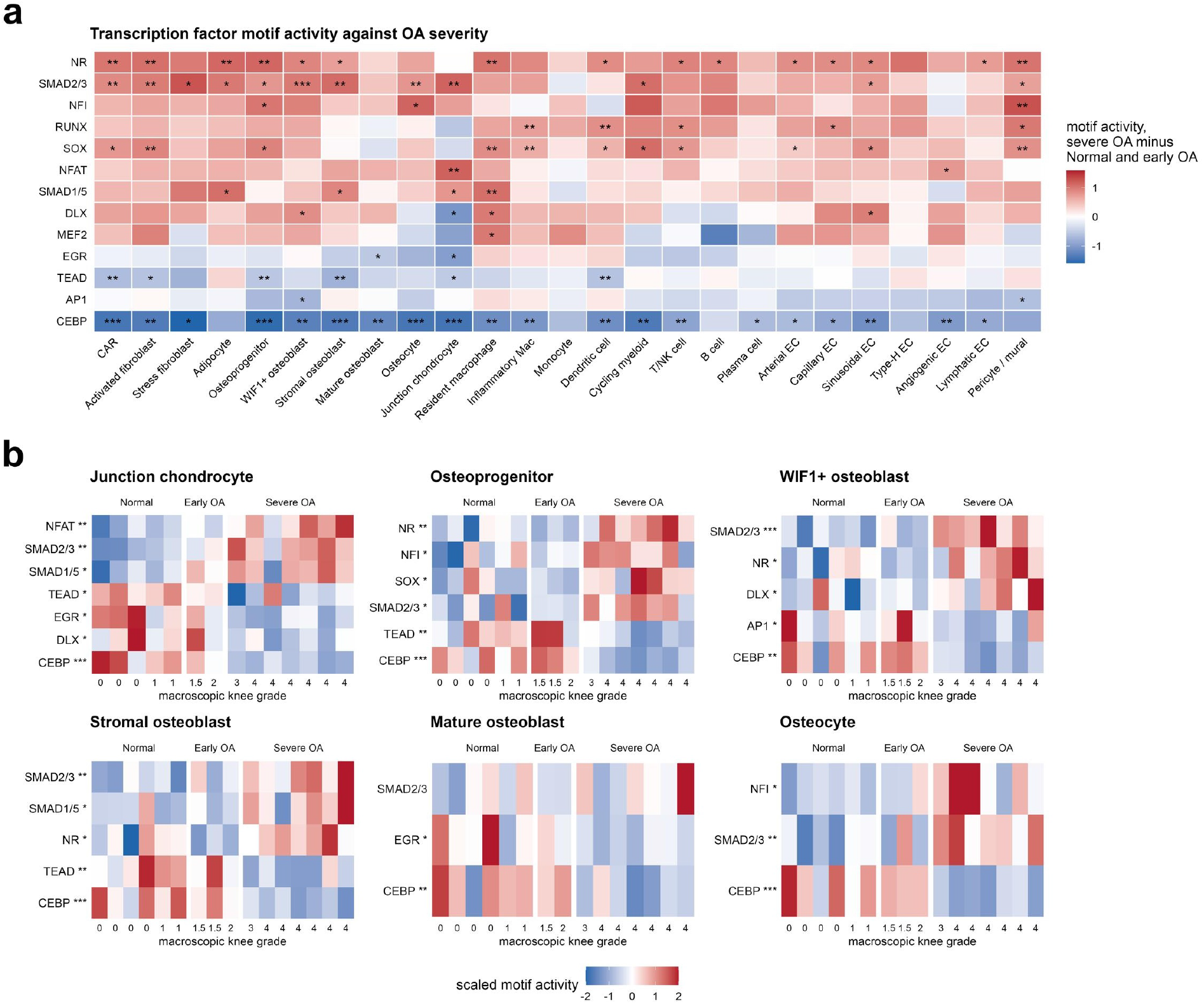
The chromatin response to severity in single-nucleus ATAC. (a) Transcription-factor family motif activity against OA severity across the 25 cell states that clear the donor threshold. Motif activity is the depth-normalized fraction of a state’s ATAC fragments falling in the motif’s top-scoring peaks (cisTarget scoring of the consensus peak set); member motifs are z-scored across donors and averaged into the family score. Tiles show the difference between severe OA and Normal and early OA donors; stars mark nominal Wilcoxon p (\**p* < 0.05, \*\**p* < 0.01, \*\*\**p* < 0.001), the convention of the composition test in Figure 7a; BH-adjusted values are in Supplementary Table 12; donor-by-state groups with at least 10 nuclei and states with at least 10 such donors. (b) The same family scores donor by donor for the junction chondrocyte and the osteogenic states from the osteoprogenitor to the osteocyte, columns ordered by macroscopic knee grade and grouped by stage. Details in Methods.

### CellChat analysis identifies the osteocyte as a sender of TGF-β and a receiver of *FN1*, and chondrocyte FGF directed at the osteoprogenitor

To identify the signals exchanged between the junction chondrocyte and the osteogenic lineage, we analyzed cell-cell communication with CellChat, first ranking every inferred pathway by the communication probability sent and received by the osteocyte (**Fig. 6a, b**). On the sending side, TGF-β, whose role in subchondral bone pathology is established,^6,8^ has the osteocyte as the tissue’s dominant source: in Normal and early OA the signal is directed mainly at the adipocyte and the resident macrophage, whereas in severe OA the TGF-β signal from the osteocyte to the junction chondrocyte rises sharply (the junction chondrocyte receives essentially no TGF-β from any state in Normal or early OA, and in severe OA nearly all of it comes from the osteocyte), and the osteocyte’s total TGF-β output increases with stage (**Fig. 6a, c**). We then asked which ligands the osteocyte itself receives. The largest inputs are matrix ligands for *CD44* (collagen, *FN1*, laminin, *SPP1*); among them *FN1* shows the largest gain from Normal to severe OA, and the *FN1* communication received by the osteocyte intensified from Normal through early to severe OA, centered on *CD44* (**Fig. 6b, d**, Supplementary Tables 9-11). *FN1* reaches the osteocyte from several senders: the junction chondrocyte’s share of this input rose from 4% in Normal to 14% in early OA and 33% in severe OA, where it was the largest single sender, and the osteoprogenitor, the *WIF1*+ osteoblast, the CAR cell and the adipocyte supplied most of the remainder (Supplementary Tables 9-11). A third, smaller axis runs from the junction chondrocyte to the osteoprogenitor and the osteoblasts: FGF signaling received by the osteogenic lineage comes from the marrow stroma in Normal and early OA, and in severe OA a chondrocyte-derived component (*FGF1* and *FGF2* through *FGFR1*) appears and accounts for a third of the FGF input (**Fig. 6e**). Two chondrocyte-derived inputs to the bone therefore emerged, FGF directed at the osteoprogenitor and the osteoblasts and *FN1* directed at the osteocyte, alongside the osteocyte’s TGF-β directed at the junction chondrocyte.

**Figure 6.**
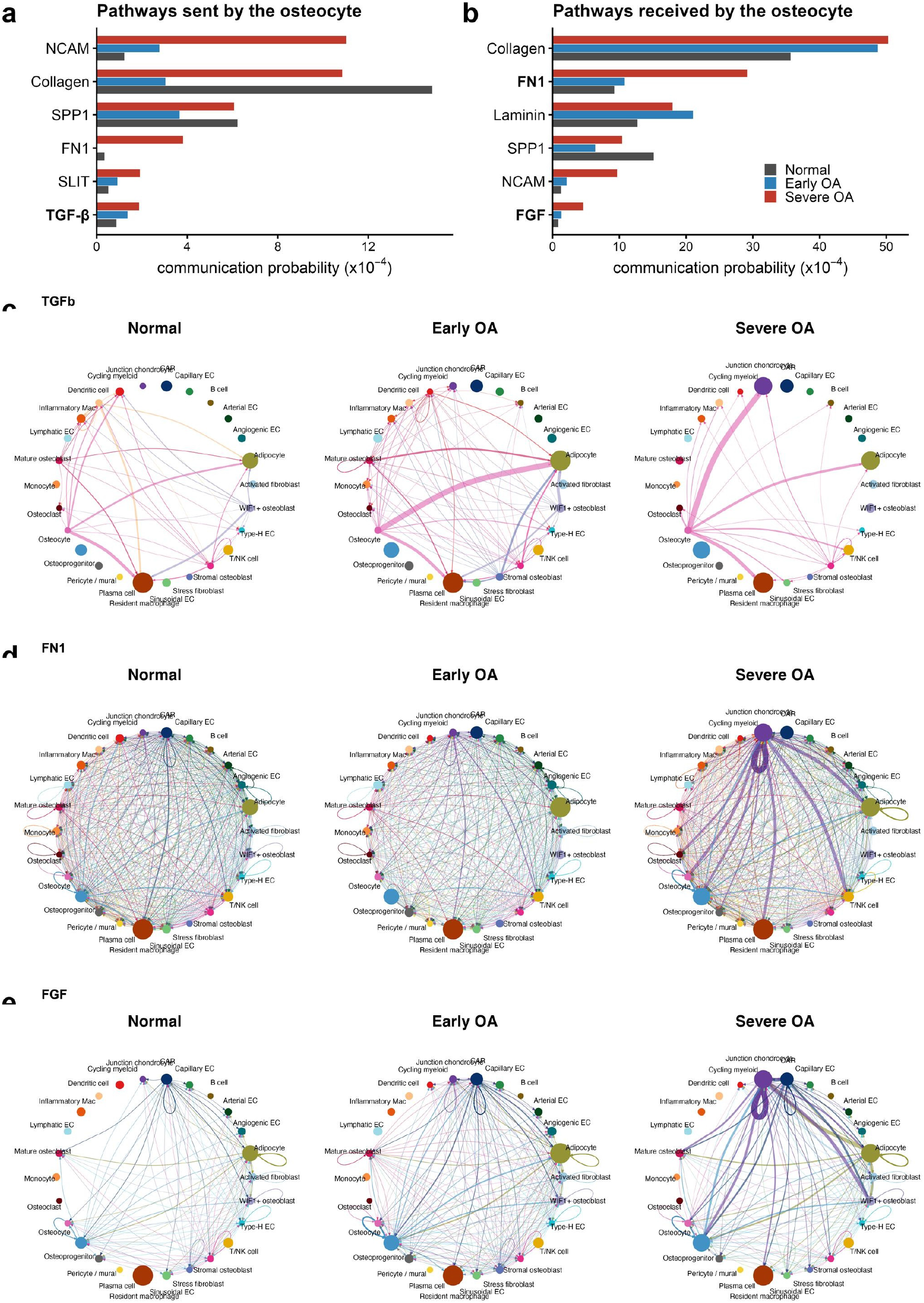
Cell-cell communication centered on the osteocyte. (a, b) The six pathways with the largest inferred communication probability sent by the osteocyte (a) and received by the osteocyte (b) in severe OA, summed over all partner states and shown per stage; the homophilic adhesion molecule PTPRM is excluded. (c) TGF-β pathway communication between the 26 cell states, one circle per stage. Edge width is the inferred communication probability; the three panels share one edge-weight maximum, and node size is the number of nuclei. (d) The same display for the FN1 pathway. (e) The same display for the FGF pathway. Details in Methods.

The osteoprogenitor and osteocyte proportions increase with OA severity We then asked whether the cell composition of the tissue changes with severity. Testing each state’s share of the donor’s nuclei against continuous grade, three states gained share significantly: the junction chondrocyte (*p* = 0.007), the osteoprogenitor (*p* = 0.003) and the osteocyte (rho = +0.54, *p* = 0.036; Fig. 7a, b; Supplementary Fig. 7, Supplementary Table 13). Within the osteocyte the gain is carried by the early osteocyte state (rho = +0.62, *p* = 0.014) rather than the mature osteocyte (rho = +0.31, *p* = 0.24; Supplementary Fig. 7c, d). To place this expansion on the differentiation axis, we drew the trajectory with Slingshot. Based on the evidence that osteoblasts differentiate from LepR-positive CAR cells,^31^ we ran the analysis with the CAR cell as the start point; the trajectory runs from the CAR cell through the osteoprogenitor and the osteoblast states to the osteocyte, and the canonical markers follow the expected pseudotime order, from *CXCL12* and *LEPR* through *RUNX2* and *SP7* to *DMP1*, *MEPE*, *PHEX* and *SOST* (**Fig. 7c, d**). The larger share of the entry state (the osteoprogenitor) and of the terminal state (the osteocyte) both rise as shares of the sampled nuclei as OA advances; the rate of differentiation is not measured here. The chromatin of the osteoprogenitor carries the corresponding signature: the motifs of the osteoblast-determining factors *SP7* (rho = 0.63, *p* = 0.008) and *DLX5* (rho = 0.63, *p* = 0.009), with *DLX3*, *SOX9* and *NFIC*, gain activity with grade, while the adipogenic *CEBPA* and *CEBPB* motifs and *ATF4* lose it (rho = -0.75, *p* < 0.001; **Fig. 7e**). In the junction chondrocyte *FGF1* and *FGF2* rise with grade (padj < 0.001 for both; Fig. 7f), and 23 of 113 ATAC peaks at *FGF1* and 12 of 46 at *FGF2* gain accessibility with grade in the same nuclei, 20 and 10 of them correlated with the gene’s RNA across donors, with none lost (Fig. 7g, Supplementary Table 14), whereas *FGFR1* and *FGFR2* on the osteoprogenitor do not change (padj = 0.56 and 0.47; Supplementary Table 6).

**Figure 7.**
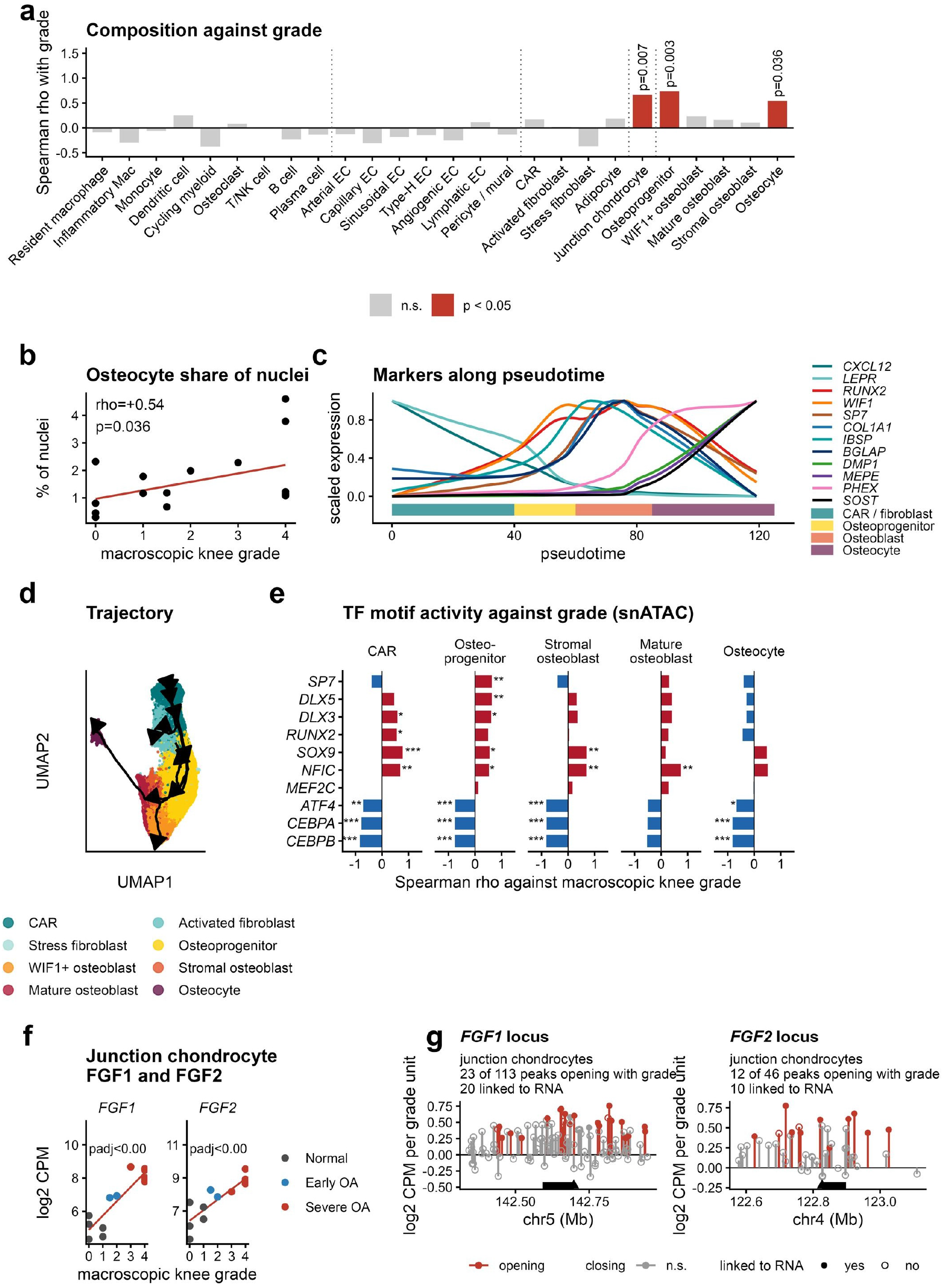
Composition and differentiation into the osteocyte. (a) Spearman correlation of each state’s abundance (centered log-ratio of the donor proportion) with continuous grade; significance by donor-label permutation, states with *p* < 0.05 in red. (b) Osteocyte share of nuclei per donor against grade. (c) Marker expression along pseudotime (loess fits, scaled per gene); the bar beneath the axis marks the state occupying each pseudotime interval. (d) Slingshot trajectory over the eight stromal-osteogenic states with the CAR cell as start cluster, on the integrated embedding; running the analysis without a start cluster returns the same ordering. (e) Motif activity of individual osteogenic and adipogenic transcription factors against continuous macroscopic knee grade in the states of the osteogenic lineage (snATAC); bars are Spearman rho across donors, stars mark nominal p (\**p* < 0.05, \*\**p* < 0.01, \*\*\**p* < 0.001) from an exploratory scan of 562 factors, and motifs of related factors (*SP7* and the DLX factors; *CEBPA* and *CEBPB*) share sequence and are not independent. (f) *FGF1* and *FGF2* in the junction chondrocyte pseudobulk against grade; each point is a donor, lines are least-squares fits, and adjusted *p* values are from the per-state model of Figure 4. (g) Consensus ATAC peaks within 250 kb of *FGF1* and *FGF2* in the junction chondrocyte (13 donors): slope of log2 CPM per grade unit; red, opening with grade (nominal *p* < 0.05); filled circles, accessibility correlated with the gene’s RNA across donors (rho ≥ 0.5, *p* < 0.05); black bar, gene body; triangle, transcription start site. Details in Methods.

### Osteocyte SMAD2/3 activity accompanies a matrix-remodeling program, with chondrocyte-derived FN1 as a candidate input

Finally, we analyzed the context of the osteocyte’s SMAD2/3 program. Osteocyte SMAD2/3 motif activity rises with grade (rho = 0.70, p = 0.003; Fig. 8a). In the osteocyte, two groups of ECM-remodeling genes that include candidate SMAD2/3 targets rise with grade (Fig. 8b): the TGF-β-responsive matrix genes *CILP*, a direct SMAD3 target,^32^ the TGF-β-binding proteoglycan *FMOD*^33^ (log2 fold change +1.71 per grade unit for *CILP*, *p* < 0.001; *FMOD p* = 0.017), *FN1* (*p* = 0.009), and the perilacunar-remodeling-associated genes^34,35^ *CA2* (*p* = 0.006), *CTSK* (*p* = 0.028) and *MMP16* (*p* = 0.034). CILP, FMOD and MMP16 correlate, across the 16 donors of the osteocyte analysis, with the osteocyte’s SMAD2/3 motif activity (rho = 0.51, p = 0.043; 0.53, p = 0.035; 0.50, p = 0.049; Fig. 8c). Along the differentiation trajectory, the FN1 receptors CD44 and ITGAV rise at the osteoblast-to-osteocyte transition, CD44 reaching 8.8-fold in the osteocyte zone of pseudotime (Fig. 8d). FN1 in the junction chondrocyte rises progressively with grade (padj < 0.001), whereas its receptors on the osteocyte do not change (padj = 0.88 and 0.73; Fig. 8e), and at the FN1 locus of the junction chondrocyte 11 of 54 peaks gain accessibility with grade, 8 of them correlated with FN1 RNA across donors, and none lose it (Fig. 8g). SMAD2/3 motif activity averaged over the osteogenic lineage tracks junction chondrocyte FN1 across donors (rho = 0.63, p = 0.017; in the osteocyte alone rho = 0.44, p = 0.12; Fig. 8f), which identifies chondrocyte-derived FN1 as a candidate input to the osteocyte SMAD2/3 program (Fig. 8a to c). Together, osteocyte SMAD2/3 motif activity rises with OA severity along with its candidate matrix and perilacunar-remodeling target genes, and across the osteogenic lineage it covaries with FN1 expression in the junction chondrocyte.

**Figure 8.**
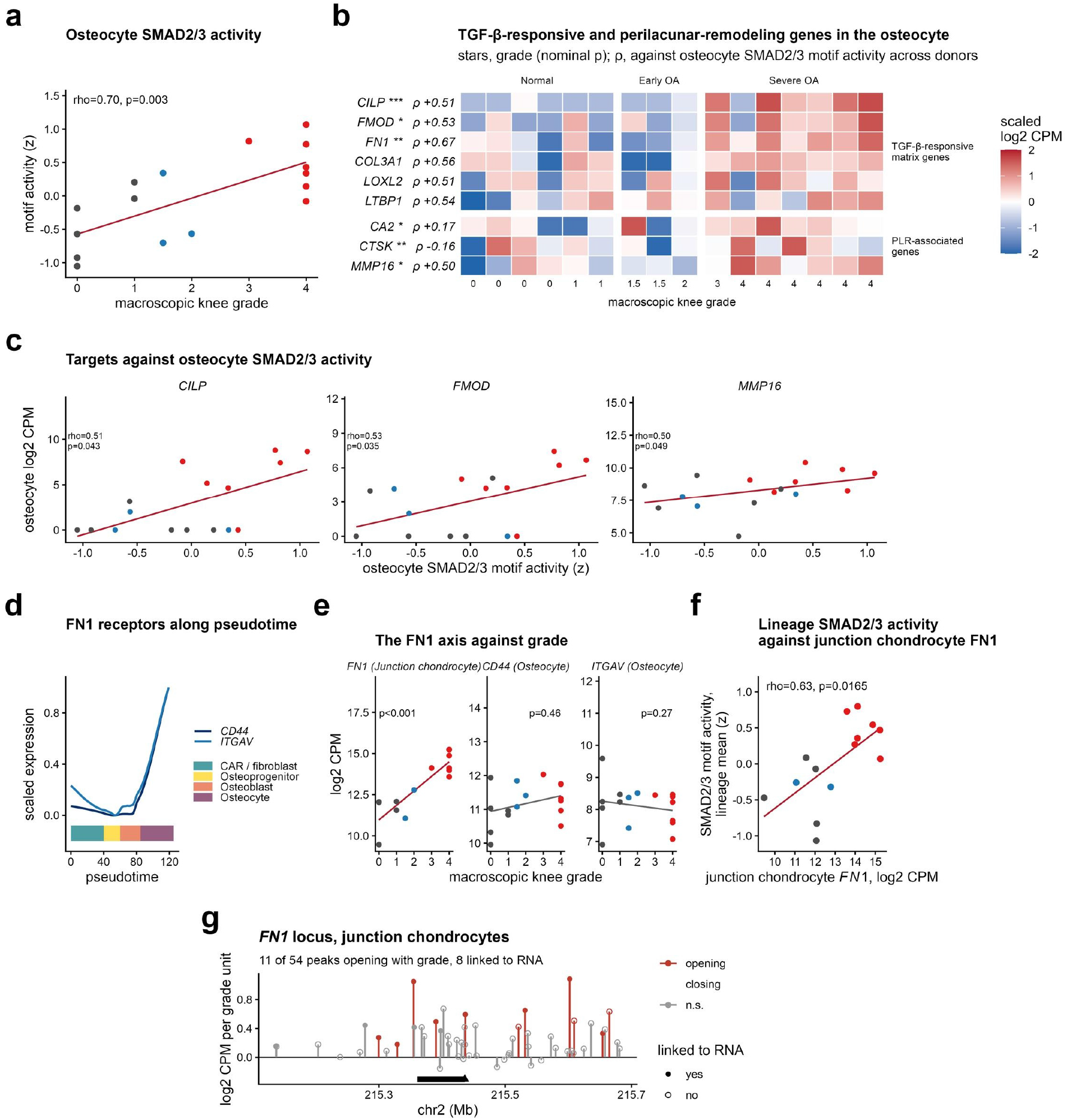
Osteocyte SMAD2/3 activity and the matrix-remodeling program, with chondrocyte-derived FN1 as a candidate input. (a) Osteocyte SMAD2/3 motif activity (family score as in Figure 5) against continuous macroscopic knee grade, one point per donor, over the same 16 donors as panels b and c; Spearman rho and nominal p. (b) Two groups of matrix-remodeling genes that include candidate SMAD2/3 targets, in the osteocyte pseudobulk over the 16 donors of the per-state model, one column per donor ordered by grade: TGF-β-responsive matrix genes (CILP, FMOD, FN1, COL3A1, LOXL2, LTBP1) and perilacunar-remodeling-associated genes (CA2, CTSK, MMP16); expression scaled per gene, stars from the per-state grade model (nominal p), and rho against osteocyte SMAD2/3 motif activity across donors. (c) CILP, FMOD and MMP16 in the osteocyte pseudobulk against the osteocyte’s SMAD2/3 motif activity, over the same 16 donors. (d) The FN1 receptors CD44 and ITGAV along the differentiation pseudotime of Figure 7 (loess fits, scaled per gene); the bar beneath the axis marks the state occupying each interval. (e) The FN1 axis against grade: FN1 in the junction chondrocyte pseudobulk and the two receptors in the osteocyte pseudobulk; each point is a donor, lines are least-squares fits, and the nominal p values of the per-state model of Figure 4 are shown; the adjusted values are given in the Results. (f) SMAD2/3 motif activity (family score as in Figure 5) averaged over the osteogenic-lineage states against FN1 expression in the junction chondrocyte pseudobulk, one point per donor (14 donors). (g) Consensus ATAC peaks within 250 kb of FN1 in the junction chondrocyte, displayed as in Figure 7g. Panels a to c are the osteocyte result; panels d to g are exploratory and identify chondrocyte-derived FN1 as a candidate input to it. Spearman rho and nominal p throughout. Details in Methods.

## Discussion

In this study we profiled human knee subchondral bone across the full severity spectrum of OA, combining bulk RNA sequencing of 89 donors with single-nucleus multiome sequencing of 16 donors, and anchored the molecular findings in the histology of the same knees. Three findings define the study. First, with advancing OA, osteoprogenitors and osteocytes occupy a larger share of the sampled nuclei, the osteocyte gain carried by the immature state, and the chondrocytes at the osteochondral junction, a matrix-modifying population whose NFAT and SMAD2/3 motifs open with disease, raise their FGF output toward the osteoprogenitor. Second, chondrocyte-derived *FN1* signaling toward the bone intensifies, a candidate link between the two tissues. Third, in step with the chondrocyte-derived *FN1* supply, SMAD2/3 motif activity rises on chromatin across the chondro-osteogenic axis, and in the osteocyte matrix and perilacunar-remodeling-associated genes are induced.

Three features give this resource its strength. First, single-nucleus analysis is what makes the osteocyte recoverable at all: the commanding cell of bone, the principal mechanosensor, the major source of RANKL and the maintainer of bone quality through perilacunar/canalicular remodeling,^34,36–40^ is embedded in mineralized matrix; none of the five published single-cell datasets of human subchondral or femoral-head bone, all enzymatic preparations of arthroplasty tissue, has recovered it,^20–24^ osteocyte-inclusive single-nucleus data exist only for the mouse,^25^ and it is the osteocyte transcriptome that carries the genetic signal for OA and osteoporosis in the mouse.^41^ Second, to our knowledge this is the first single-nucleus multiome, including snATAC, applied to adult human subchondral bone. Paired single-nucleus transcriptome and chromatin profiling of human skeletal tissue has been reported for the embryonic limb, joint and cranium,^42^ and for the adult osteogenic lineage only in the mouse,^24,26^ so transcription and transcription-factor activity are measured in the same nuclei, osteocytes included. Third, the cohort of 89 bulk and 16 multiome donors including truly normal and early-OA knees is the largest reported for human subchondral bone, and turns disease severity into a continuous axis rather than an end-stage contrast.^16,17,20^

The first axis of the disease response is quantitative: transcriptomic changes consistent with bone gain. The junction chondrocyte, the osteoprogenitor and the osteocyte take a larger share of the nuclei with grade, and the trajectory from the CAR cell through the osteoprogenitor and the osteoblast states to the osteocyte agrees with mouse lineage tracing.^31,43^ Because the CAR pool itself does not shrink while the shares of the osteoprogenitor and the osteocyte grow, the pattern is consistent with sustained entry into the lineage rather than exhaustion of the reservoir, although the rate of differentiation is not measured directly here; the chromatin of the osteoprogenitor, in which the *SP7* and *DLX5* motifs open and the adipogenic CEBP motifs close with grade, points the same way. A larger osteocyte share of the sampled nuclei is not a higher osteocyte density in bone. Histologically, osteocyte lacunar density rises with cartilage degeneration in knee and hip OA while osteocyte viability falls and empty lacunae increase,^44,45^ and the sclerotic zones of OA bone carry osteocytes of immature phenotype.^46^ The increased fraction may reflect changes in the relative representation of bone and marrow compartments, newly formed osteocytes, or differences in nuclear recovery across disease stages. Its concentration in the early osteocyte state is consistent with reports of immature osteocytes in sclerotic bone, but neither absolute osteocyte density nor survival of the existing network was measured here. The thickening of the subchondral bone plate in advanced OA^3^ is consistent with this expansion, and the activation of bone-matrix and ossification programs, which builds with grade, captures at the transcriptional level the transition into the late, sclerotic phase of the biphasic remodeling of OA bone.^3^ Among individual genes, *TNFRSF11B*, encoding the anti-resorptive decoy receptor osteoprotegerin (OPG), rose with grade in the osteocyte and the osteoprogenitor while *TNFSF11*, encoding RANKL, was unchanged;^47,48^ an abnormal OPG/RANKL balance has also been reported in OA subchondral osteoblasts.^49^ OPG is a protective factor against resorption, and its administration reduces cartilage degeneration and pain in animal models.^50,51^ Because the osteocyte is itself the major source of RANKL,^38,39^ the rise of OPG in the osteocyte appears as a compensatory response restraining the remodeling activated by advancing OA; a resulting restraint of resorption, which we did not measure, would fit the formation-dominant remodeling and sclerosis of late disease.

To ask what drives this entry into the osteogenic lineage, we considered the FGF signal from the junction chondrocyte, because three features of the osteoprogenitor in our data match FGF functions in marrow stromal progenitors. The osteoprogenitor gains share with grade (Fig. 7a), its chromatin opens the motifs of the osteoblast-determining factors *SP7* and *DLX5* and closes those of the adipogenic CEBP factors (Fig. 7e), and in severe OA about a third of the FGF it receives comes from the junction chondrocyte, through FGF1 and FGF2 acting on FGFR1 (Fig. 6e); in the junction chondrocyte *FGF1* and *FGF2* rise with grade, while *FGFR1* and *FGFR2* on the osteoprogenitor stay constant (Fig. 7f, Supplementary Table 6). Mice lacking Fgf2 lose trabecular bone through reduced bone formation,^52^ and their marrow stromal cells form fewer alkaline phosphatase-positive, mineralizing colonies and more adipocytes, with induction of PPARγ2, a shift that exogenous FGF2 reverses.^53^ FGFR1 acts at successive steps of the lineage: its loss in osteo-chondroprogenitors delays osteoblast differentiation, whereas its loss in differentiated osteoblasts accelerates their maturation and mineralization,^54^ so FGF signaling through FGFR1 promotes the commitment of progenitors while restraining the terminal maturation of osteoblasts.^55^ A rising ligand supply on constant receptors, an osteoprogenitor whose chromatin turns from the adipogenic toward the osteogenic program, and a larger share of the entry state are therefore three readouts of one process. The cartilage is also a potential source of the ligand: articular cartilage stores FGF2 in its pericellular matrix and releases it on mechanical injury, where it activates ERK through FGFR1,^56^ OA chondrocytes drive subchondral osteoblasts toward an osteogenic phenotype through ERK signaling in co-culture,^57^ and within the cartilage FGF2 suppresses ADAMTS5, its loss accelerating murine OA.^58^ We therefore propose that junction chondrocyte-derived FGF may influence osteogenic commitment as OA advances, potentially coupling the response of deep cartilage to remodeling of the underlying bone. This mechanism requires functional testing.

To define how the cartilage facing the bone changes with OA, we examined the junction chondrocyte, the cells of the deep and calcified cartilage that remain attached to the subchondral bone when the overlying cartilage is harvested and are therefore rarely profiled. Their most specific transcripts (*ACAN*, *CILP*, *COMP*, *PRELP*, *ECRG4*, *TNFRSF11B*) mark the deep and calcified zones, and with grade they intensify a matrix program rather than entering hypertrophy: collagen fibril organization, cartilage development and ossification rise, together with *FN1*, *FGF1*, *FGF2*, *HTRA1*, *COMP* and *TNFSF11*, while *COL10A1*, *IBSP* and *RUNX2* stay flat (Fig. 4e, i, Supplementary Table 14). Their chromatin moves in the same direction. The SMAD2/3 and NFAT motifs open, both in the motif activity of the state and among the 29,870 peaks that gain accessibility with grade, whereas the CEBP, EGR and KLF/SP motifs close (Fig. 5a, b, Supplementary Table 14). Each of these regulators has a defined role in articular cartilage. TGF-β/SMAD3 signaling represses the terminal hypertrophic differentiation of chondrocytes, and its loss produces OA with type X collagen-expressing chondrocytes;^59^ NFAT1 maintains the adult articular chondrocyte, and its loss causes OA through combined catabolic and anabolic failure;^60^ and C/EBPβ cooperates with RUNX2 to induce MMP13 and degrade cartilage during endochondral ossification and OA.^61^ The junction chondrocyte of the OA knee therefore opens the programs that restrain hypertrophy and closes the one that drives it, a matrix-producing response of the deep cartilage rather than its terminal differentiation.

This state of the junction chondrocyte bears directly on the two structural changes of the osteochondral junction in OA, the multiplication of vascular channels and the thickening of the calcified layer.^12,13,62^ Channels form when osteoclasts and vessels from the subchondral marrow breach the bone plate and the calcified cartilage.^12,13^ The junction chondrocyte raises *TNFSF11*, encoding RANKL (log2 fold change +1.10 per grade unit; Fig. 4i), and in the growth plate it is the RANKL of the matrix-embedded hypertrophic chondrocyte that controls the resorption of mineralized cartilage;^39^ the junction chondrocyte is therefore positioned to open channels from the cartilage side, while the osteocyte and the osteoprogenitor beneath it raise OPG (Fig. 4f, g). Its FGF1 and FGF2 then reach the osteoprogenitors that accompany type H vessels into the marrow spaces,^7^ biasing them toward bone as discussed above. The calcified cartilage, for its part, thickens in OA by endochondral advance of the tidemark and replacement of calcified cartilage by bone, which contributes to subchondral sclerosis,^62^ and the hydraulic conductance across the cartilage and the subchondral plate rises as the disease progresses.^63^ In our data the chondrocyte at the junction does not take up the hypertrophic program; the advance is accompanied instead by the matrix and ossification programs that rise in this chondrocyte and by the expanding osteogenic lineage beneath it. The molecular state of the junction chondrocyte is thus positioned to contribute to both structural changes, RANKL and FGF that open and fill the channels and a matrix program that thickens the calcified layer without terminal hypertrophy, and identifies this population as a candidate mediator of reciprocal cartilage–bone remodeling in OA.

The same bulk RNA-seq data also reveal a change in transcript structure that is not visible when expression is summed over each gene. Across the 89 knees, isoform usage converged on each gene’s principal transcript as grade advanced (Fig. 2d, e), and one of the shifts concerns the oncostatin M receptor. *OSMR* is transcribed as the membrane receptor (OSMR-201) and as a transcript that ends within intron 7 and encodes a 342-residue protein without the transmembrane segment (OSMR-202); the secreted product of this transcript, soluble OSMR, binds oncostatin M and interleukin-31 and neutralizes both.^64^ With grade the soluble transcript lost 4.8 points of *OSMR* usage per grade unit and the membrane transcript gained 6.1, while total *OSMR* expression did not change (Fig. 2g). Oncostatin M, produced by neutrophils and activated macrophages, drives the osteoblast differentiation and mineralization of marrow stromal cells through OSMR, gp130 and STAT3,^65^ and signaling through OSMR in osteoblasts induces RANKL; mice lacking Osmr are osteopetrotic.^66^ The shift in OSMR isoform usage is consistent with altered potential responsiveness to oncostatin M, although receptor protein abundance and signaling activity were not measured.

The second axis is qualitative, and it crosses tissues. Junction chondrocyte *FN1* expression rises linearly with OA severity. Fibronectin in OA has so far been a damage signal completed within cartilage;^67,68^ in our data the junction chondrocyte supply of the ligand rises with severity. The osteochondral channels connecting cartilage and bone multiply with disease stage,^12^ expanding the physical route for the ligand, and the receiving side is defined: roughly 80% of the FN1 input onto the osteocyte is carried by CD44, itself an osteocyte marker,^69^ and *CD44* and *ITGAV* rise precisely at the osteoblast-to-osteocyte transition. The cell prepared to receive the increased *FN1* is therefore the osteocyte. From this a hypothesis follows. Fibronectin fibrils are required for the deposition of latent TGF-β binding protein 1 into the matrix,^70^ *CD44* and αv integrins associate with the TGF-β receptor complex and potentiate its signaling, shown for hyaluronan-engaged *CD44* and for αvβ3,^71–73^ and the osteocyte is the tissue’s dominant *TGFB2* source; *FN1* arriving from the cartilage may therefore raise the osteocyte’s sensitivity to its own TGF-β, through availability, sensitivity or both. Osteocyte SMAD2/3 activation in human advanced OA subchondral bone has been reported at the protein level, where osteocyte phospho-SMAD2/3 correlates with bone mineral density and formation markers;^11^ our chromatin data are consistent with it and show that the gain is shared across the chondro-osteogenic axis. Across the osteogenic lineage the mean SMAD2/3 motif activity covaries with junction chondrocyte FN1, while the association in the osteocyte alone is not resolved (rho = 0.44, p = 0.12), and among the candidate SMAD2/3 targets, CILP, a direct SMAD3 target,28 the TGF-β-binding proteoglycan FMOD29 and the perilacunar-remodeling-associated genes,30,31 FMOD rises with the osteocyte’s own SMAD2/3 activity. These observations nominate FN1 as a candidate input and do not establish an FN1-to-osteocyte mechanism. Other grade-responsive junction chondrocyte ligands rise in parallel, among them *COMP*, a TGF-β-binding matrix protein that enhances TGF-β activity,^74^ and *INHBA*, and at the junction *FN1* rises within the same matrix-modifying program as *FGF1* and *FGF2* (Fig. 4i), so the between-donor correlations do not single out *FN1*, and we treat it as the leading candidate. The direction in which the induced program changes bone quality is not settled: in the mouse, loss of osteocyte TGF-β signaling or of *MMP13* suppresses perilacunar remodeling and degrades bone quality and the joint,^34,35^ so both suppression and dysregulated induction of the program are compatible with impaired matrix maintenance. What our data show is that the program is induced in the osteocyte with advancing severity; whether that response is maladaptive or compensatory cannot be resolved here. On this view, TGF-β-directed therapy for subchondral bone^6,8^ would act on the osteocyte, although whether its effect there is beneficial remains to be determined.

The two directions of signaling across the osteochondral junction may form one circuit, and the chondrocytes at the junction define it. The osteocyte’s TGF-β output in severe OA is directed above all at the junction chondrocyte: in the communication analysis the junction chondrocyte receives essentially no TGF-β from any state in Normal or early OA, and in severe OA nearly all of its TGF-β input comes from the osteocyte (**Fig. 6c**), with the multiplying osteochondral channels^12^ as a route. On the chromatin of the junction chondrocyte, SMAD2/3 motif activity shows the largest significant gain of the chondro-osteogenic axis (+1.16, p = 0.004; Fig. 5a), and in its own peaks the SMAD2/3 and NFAT motifs are enriched among the regions that open with grade, including those at the *FGF1*, *FGF2* and *FN1* loci (Fig. 7g, 8g, Supplementary Table 14). Signaling through ALK5 and SMAD2/3 restrains chondrocytes from terminal hypertrophy,^59,75^ so a plausible circuit runs from osteocyte TGF-β to the junction chondrocyte, whose SMAD2/3 motifs open with grade while its hypertrophy genes stay flat, from its FGF to the osteoprogenitor and osteoblast commitment, and from the resulting bone formation to the cartilage again; it would give the finding that excess TGF-β in subchondral bone drives cartilage degeneration in the mouse^6^ a cellular reading in human tissue. The opening of SMAD2/3 without a hypertrophic response places the junction chondrocyte on the ALK5 side of the shift of TGF-β signaling from ALK5 toward ALK1 described in aged and OA cartilage.^75^

This study has several limitations. The cohort is cross-sectional, so the causal order of the model, including whether FN1 amplifies SMAD2/3 signaling and whether junction chondrocyte FGF drives the commitment of the osteoprogenitor, is an inference from between-donor covariation, and functional validation remains for future work. CellChat infers signaling from ligand and receptor expression and does not measure transmission at the protein level. In particular, transcript-based inference cannot distinguish intact fibronectin from the fibronectin fragments that accumulate in the OA joint and drive chondrocyte catabolism through the α5β1 integrin and Toll-like receptor 2.^67,76,77^ The CellChat fibronectin pathway models the intact ligand against six receptor pairs and does not include Toll-like receptors; in our data the bone states express little *ITGA5*, and *TLR4* falls with grade in the osteocyte and the osteoprogenitor (Supplementary Table 6), so the inferred input onto the osteocyte rests on *CD44* and αv integrins, which bind the heparin-binding and RGD domains present in intact fibronectin and in its larger fragments. Which form of fibronectin reaches the bone needs to be determined. Motif activity summarizes accessibility over many peaks per factor family and does not resolve individual loci, and lacunar-canalicular morphometry was not part of this study.

In conclusion, human OA subchondral bone shows two coupled responses centered on the osteocyte and the osteochondral junction: a quantitative response, expansion of the osteogenic lineage, for which FGF from the junction chondrocyte is a candidate driver, and a qualitative response, a rise of SMAD2/3 activity across the chondro-osteogenic axis with the osteocyte’s matrix-maintenance program, for which chondrocyte-derived FN1 is a candidate input (Fig. 9). These findings identify osteocytes and the osteochondral junction as priorities for mechanistic validation and therapeutic investigation in OA.

**Figure 9.**
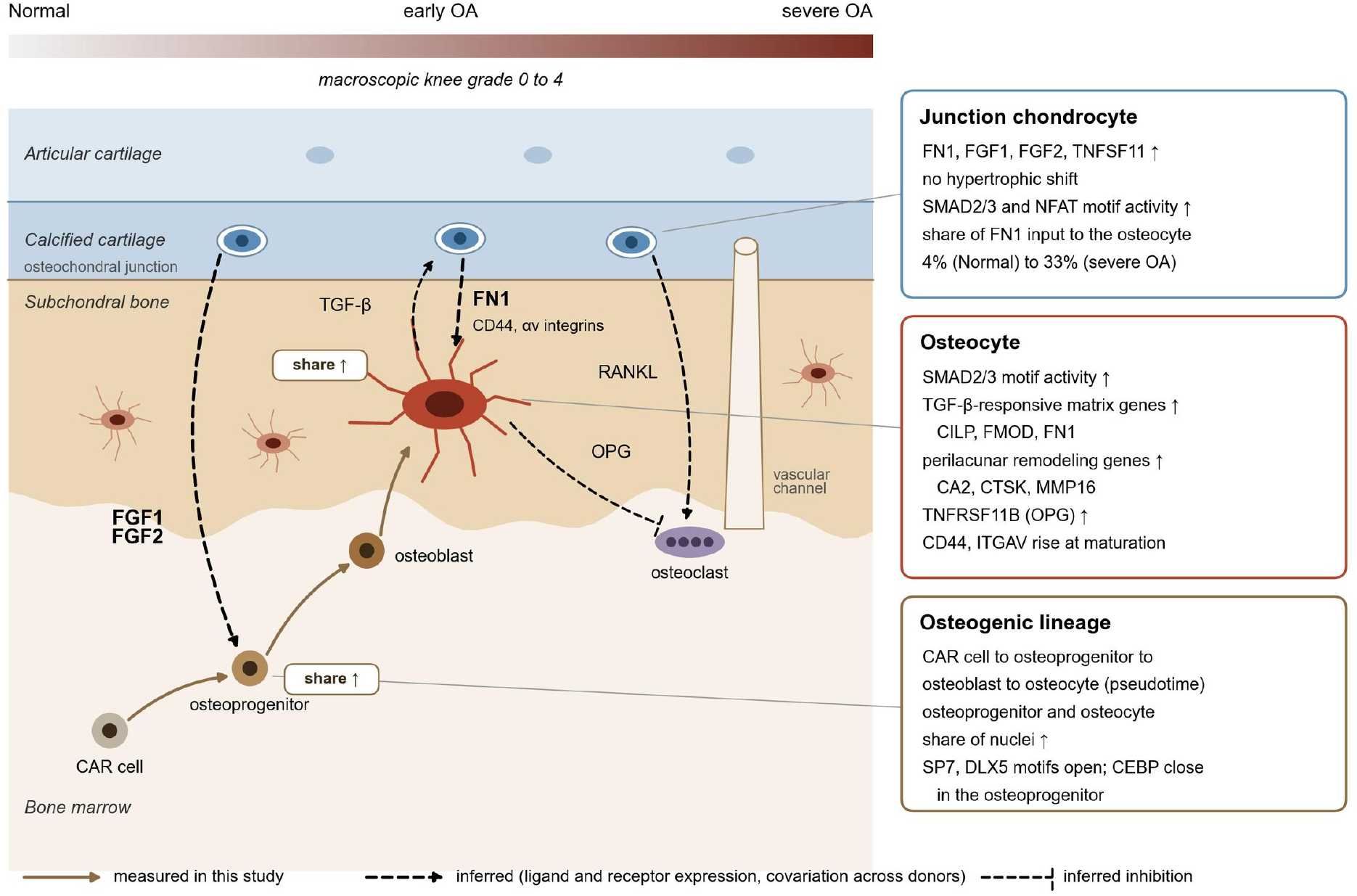
Summary: the osteocyte at the center of the subchondral bone response to OA. With advancing grade, the osteogenic lineage runs from the CXCL12-abundant reticular (CAR) cell through the osteoprogenitor and osteoblast states to the osteocyte, and the osteoprogenitor and the osteocyte take a larger share of the nuclei (Fig. 7). Chondrocytes at the osteochondral junction raise FN1, FGF1, FGF2 and TNFSF11 (RANKL) without a hypertrophic shift (Fig. 4); FGF1 and FGF2 are inferred to reach the osteoprogenitor and FN1 to reach the osteocyte through CD44 and αv integrins (Figs. 6 and 8). In the osteocyte, SMAD2/3 motif activity rises with grade together with TGF-β-responsive matrix genes (CILP, FMOD, FN1) and perilacunar-remodeling genes (CA2, CTSK, MMP16), and TNFRSF11B (OPG) rises (Figs. 4, 5 and 8). The TGF-β signal from the osteocyte to the junction chondrocyte rises in severe OA (Fig. 6). RANKL from the junction chondrocyte and OPG from the osteocyte are drawn toward the osteoclast at a vascular channel; their effect on resorption is inferred. Solid arrows, changes measured in this study; dashed arrows and the dashed bar, signals inferred from ligand and receptor expression or from covariation across donors.

## Methods

### Human subjects and tissue

All human tissues were obtained with approval by the Scripps Human Subjects Committee (protocol 20-7636). Patients undergoing total knee arthroplasty were enrolled at Scripps Clinic and provided written consent; intact knee joints from organ donors were procured by Lifesharing. Disease severity was scored macroscopically across all knee cartilage surfaces with the Outerbridge method^27^ (macroscopic knee grade, 0 to 4), used throughout as a continuous variable. Organ donors spanned grades 0 to 4 and every arthroplasty knee is grade 4; of the 51 grade 4 knees, 48 are arthroplasty specimens and 3 are organ donors. For staged displays, donors were grouped on one severity axis as Normal (grade 0 to 1), early OA (grade 1.5 to 2) and severe OA (grade 3 to 4). The bulk cohort comprises 89 knees with macroscopic grade available (20 Normal, 16 early OA, 53 severe OA) and the single-nucleus cohort 16 donors (6, 3 and 7), spanning ages 18 to 84. Donor information is given in Supplementary Tables 1 and 2.

### Osteochondral histopathology

Osteochondral specimens were taken from the weight-bearing region of the medial femoral condyle (organ donors) or from the surgical resection from the same area (arthroplasty), fixed in Z-fix (Anatech), decalcified in 10% EDTA for 3 to 16 days, embedded in paraffin, sectioned at 5 µm and stained with Safranin-O/fast green. Slides were scanned on an Aperio AT2 whole-slide scanner (Leica Biosystems) and analyzed in QuPath v0.5.1. Cartilage changes were assessed with the OARSI scoring system^28^ and bone changes with the OABS.^29^

### RNA isolation

To minimize tissue degradation, collection and processing followed the published protocols (protocols.io; Data availability). Total RNA was isolated directly from frozen subchondral bone pulverized under liquid nitrogen.

### Nuclei isolation

Nuclei were isolated from frozen subchondral bone stored in liquid nitrogen, following the published protocol (https://dx.doi.org/10.17504/protocols.io.j8nlk8mrdl5r/v1). Before loading, 200 mg of frozen tissue was pulverized under liquid nitrogen with a mortar and pestle. The tissue fragments were transferred to a 50 mL tube with 2 mL of lysis buffer and incubated for 20 min at 4 °C, after which 2 mL of wash buffer was added. The suspension was passed through 40 µm, 20 µm and 10 µm strainers to remove tissue and cellular debris, and the flow-through was collected on ice and transferred to 5 mL LoBind tubes. After centrifugation at 500 × g for 5 min at 4 °C, the supernatant was removed and the nuclei pellet was resuspended in 400 µL of PBS containing 0.03 mg/mL recombinant albumin and 1 U/µL RNase inhibitor. To further eliminate debris, a second centrifugation was performed at 150 × g for 10 min at 4 °C, and the pellet was resuspended in the same buffer. For counting and quality assessment, 18 µL of the suspension was mixed with 2 µL of acridine orange and counted on a Countess II FL automated cell counter (AOPI staining); nuclear morphology and membrane integrity were inspected under a microscope at 40x or 60x, following the nuclear integrity guidelines of 10x Genomics. Based on the estimated concentration, the pellet was resuspended in chilled diluted nuclei buffer to 4,000 to 5,000 nuclei/µL for a final recovery of 20,000 nuclei in 5 µL.

### Library preparation and sequencing

Single-nucleus multiome libraries (gene expression and ATAC) were generated with the Chromium Next GEM Single Cell Multiome ATAC + Gene Expression Reagent Kit (10x Genomics), which profiles chromatin accessibility and gene expression from the same nucleus. Transposase first fragmented DNA in open chromatin and added adapter sequences; nuclei were then partitioned into Gel Beads-in-Emulsion, where transposed DNA (ATAC) and cDNA (gene expression) were separately barcoded. After post-GEM cleanup, primers were added for reverse transcription and amplification, and the pre-amplified DNA and cDNA were used to construct the ATAC and gene expression libraries, which were PCR-amplified. Library quality was assessed on an Agilent Bioanalyzer 2100. The gene expression and ATAC libraries were loaded onto separate P3 flow cells for paired-end sequencing on an AVITI instrument (Element Biosciences), targeting 10,000 nuclei per sample at 25,000 reads per nucleus. Bulk RNA sequencing (https://dx.doi.org/10.17504/protocols.io.6qpvr8m2zlmk/v1) used ribosomal RNA depletion and NovaSeq X sequencing at approximately 30 million read pairs per sample.

### Single-nucleus processing, clustering and annotation

Reads were processed with Cell Ranger ARC (GRCh38-2020-A, intronic reads counted). Nuclei were retained at 1,000 to 25,000 RNA counts and below 5% mitochondrial reads, giving 62,993 nuclei (23,495 Normal, 12,382 early OA, 27,116 severe OA). After annotation, 1,423 nuclei of the chondrocyte state were excluded by a transcript-based filter (see Quality filter of the chondrocyte state), leaving 61,570 nuclei (22,552 Normal, 12,220 early OA, 26,798 severe OA) for all downstream analyses. Samples were normalized with SCTransform, integrated with Seurat canonical correlation anchors and clustered by Louvain community detection at a resolution selected by silhouette width and modularity; five labels comprising 3,657 nuclei were removed as quality artifacts. Identities were assigned from a literature audit of marker genes (**Fig. 3c**). The clustering resolved two osteocyte states, of which the early osteocyte falls below the testability floor on its own; they are presented and tested as a single osteocyte cluster, giving 26 cell states. Ambient RNA was flagged but not corrected in the presented data; because lineage-restricted matrix genes are detected across lineages, cross-lineage matrix expression is not interpreted as biology in this manuscript.

### Capture-burden covariate

Single-nucleus libraries over-represent long genes, and the per-sample share of long-gene signal covaries with grade in this cohort. A capture-burden statistic (fraction of UMIs in genes with genomic span of at least 110.4 kb, restricted to the 4,022 long genes without ECM or skeletal annotation) was computed per sample and cluster and included as a covariate in all single-nucleus differential models. The bulk libraries show no length-associated signal (r = +0.003 across 15,026 genes).

### Principal component analysis of the bulk cohort

Counts were filtered to genes with at least 10 reads in at least 10 samples (25,480 genes), transformed to log2 counts per million, and the 2,000 most variable genes entered a centered, unscaled principal component analysis over the 89 donors. Stages were compared on the components by two-sided Wilcoxon rank-sum test.

### Bulk differential expression and gene set enrichment

Three hypotheses were specified before the analysis and are treated as confirmatory: the expansion of the osteocyte and of the osteoprogenitor with grade, and the rise of SMAD2/3 motif activity in the osteocyte. Each is nominally significant (osteoprogenitor expansion p = 0.0027, osteocyte expansion p = 0.036, osteocyte SMAD2/3 motif activity p = 0.003), and all three remain significant when the three are treated as one family and Holm corrected (0.008, 0.036 and 0.008). All other comparisons, including the remaining composition, motif-activity and donor-level correlation results, are exploratory, and their p values are nominal unless an adjusted value is given. Bulk counts were tested with DESeq2 under the design ∼ batch + detected genes + macroscopic knee grade (continuous), where the detected-genes covariate is the standardized number of genes detected per sample. Genes passing the expression filter of the run of record (17,366 genes) were tested; significance is Benjamini-Hochberg adjusted p below 0.05 (Supplementary Table 3). The same model was refitted with the macroscopic grade replaced by the OABS score of the medial weight-bearing subchondral bone of the same knee, available for 85 of the 89 knees. The osteoarthritis bone score (OABS) is a published histological score of subchondral bone in which seven features of bone-marrow-lesion pathology are recorded as present or absent and summed, giving 0 to 7 per region;^29^ it was read on the same sections as the OARSI cartilage score, one score per knee. The refitted model was run on the same gene universe with the Benjamini-Hochberg correction recomputed over it (Supplementary Fig. 2). Pre-ranked GSEA (GO biological process) used the ranking −log10(p) × sign(log2 fold change) with gseGO (clusterProfiler; gene sets of 10 to 500 members, BH adjustment); significant terms are given in **Supplementary Table 4**.

### Isoform usage in bulk RNA-seq

Transcript abundances were estimated from the HISAT2 alignments of the 89 knees with StringTie v1.2.3^78^ in reference-guided mode (options -e -B) against GENCODE v44,^79^ the annotation of the gene-level analysis, and imported with IsoformSwitchAnalyzeR^80^ (read length 150). Genes expressed at 1 TPM or more with more than one isoform were kept (90,378 isoforms of 14,100 genes), and the isoform fraction, the share of a gene’s expression carried by each isoform, was computed per knee. For each isoform the fraction was regressed on batch, the detected-genes covariate and continuous macroscopic knee grade; the slope, in percentage points per grade unit, is the reported effect, and the p value is that of the Spearman correlation between the fraction and grade after both were adjusted for batch and the detected-genes covariate. Isoforms with p < 0.05 and a slope of at least 1.25 points per grade unit in either direction are reported (Supplementary Table 5). Isoform classes were defined against each gene’s principal isoform, the isoform with the highest mean fraction in normal knees: alternative transcription start or 3′ end (more than 100 bp from those of the principal isoform), longer or shorter 3′ UTR (by more than 100 nt), and retained-intron, nonsense-mediated decay and non-coding transcripts according to the GENCODE biotype. Enrichment of each class among the rising and among the falling isoforms relative to all tested isoforms was tested with Fisher’s exact test. Fig. 2f shows the strongest rising and falling isoforms among those with a median fraction between 0.1 and 0.9 and a fraction between 0.02 and 0.98 in at least 70% of knees. Protein consequences were read from the GENCODE coding sequences.

### Single-nucleus differential expression and gene set enrichment

Counts were aggregated to pseudobulk per sample and cluster, and DESeq2 was fitted with the design ∼ capture burden + macroscopic knee grade (continuous). Samples entered a cluster’s model with at least 10 nuclei, and populations were tested with at least 10 such samples. The same rule is applied to the motif-activity analysis of the single-nucleus ATAC, so the per-state expression model and the motif activity of a state rest on the same donors. Two states fall below the threshold: the osteoclast with 7 donors and the early osteocyte with 9. Their results are given in Supplementary Table 6 with the donor count and are not shown as expression or motif-activity panels. The merged osteocyte population has 16 donors. For the chondrocyte state the model was fitted on the junction chondrocytes (14 donors with at least 10 nuclei). Significance is Benjamini-Hochberg adjusted *p* below 0.05 (Supplementary Table 6). The two osteocyte states were merged into one population (raw-sum pseudobulk, covariate recomputed identically; all 16 donors testable). Per-cluster pre-ranked GSEA used the same ranking and settings as the bulk analysis; significant terms for the osteogenic states are given in Supplementary Table 7. In the GSEA panels of **Fig. 4**, up to four significant terms per direction are shown, with skeletal-program terms (ossification, skeletal, cartilage, chondrocyte, ECM, biomineral) shown first; ribosomal and translation terms are treated as a library-level property of single-nucleus data. The bulk volcano plot labels, and the per-state heatmaps include, the genes of a pre-specified TGF-β and FGF axis list that reach BH-adjusted *p* < 0.05 (ligands *TGFB1*, TGFB2, TGFB3 and INHBA; receptors TGFBR1, *TGFBR2*, *TGFBR3*, *FGFR1*, FGFR2 and FGFR3; latent-complex proteins LTBP1 to LTBP4; *FGF1*, *FGF2*, FGF7, FGF9 and FGF18; and the TGF-β-responsive matrix genes *COL1A1*, *COL1A2*, *COL3A1*, *BGN*, DCN, *FN1*, *CILP*, *FMOD*, *COMP*, POSTN, SERPINE1, CCN2, TGFBI and THBS1); the per-state heatmaps otherwise show the characterizing genes of that state, the remaining rows filled by smallest adjusted *p*.

### Composition against grade

Abundance was tested per cell state as the Spearman correlation between the centered log-ratio transformed donor proportion (pseudocount 0.5 nuclei) and continuous macroscopic knee grade, across the 16 single-nucleus donors; the 1,423 nuclei excluded from the chondrocyte state were not counted. Significance was assessed by donor-label permutation (10,000 permutations, two-sided), and states with permutation p below 0.05 are reported.

### Quality filter of the chondrocyte state and chondrocyte chromatin analyses

Nuclei labeled chondrocyte in the atlas (5,402) were scored for module scores of chondrocyte (*COL2A1*, *ACAN*, *HAPLN1*, *SOX9*, *COMP*, *CILP*, *ECRG4*, *PRG4*, *CRTAC1*, *CHI3L1*), myeloid, erythroid and plasma-cell programs (Seurat AddModuleScore). A nucleus was reassigned to a non-chondrocyte program when that score exceeded 0.3 and at most 3 of the 10 chondrocyte genes were detected, and flagged as low content when at most 2 chondrocyte genes were detected. In normal knees 27% of the labeled nuclei passed (36% erythroid, 19% low content, 12% plasma cell, 7% myeloid), against 72% in early OA and 97% in severe OA (Supplementary Table 14). The 4,273 passing nuclei were integrated across donors by reciprocal PCA (Seurat v5; donors with at least 100 nuclei as separate batches, smaller donors pooled; 2,000 features, 30 dimensions, k.weight 20) and clustered (resolution 0.3). Four small subclusters that co-expressed macrophage (136 nuclei), endothelial (75), adipocyte (51) or T and NK (32) genes were removed, leaving 3,979 junction chondrocytes in five subclusters; subcluster names reflect their top markers. The 1,423 excluded nuclei were removed from the chondrocyte state for every downstream analysis (composition, per-state differential expression, CellChat and the ATAC pseudobulks); the clustering, the other 25 states and their counts are unchanged. Specificity against the other states was computed as the difference between the mean log-normalized expression in the junction chondrocytes and the maximum mean in any other state. Per-donor pseudobulk log2 CPM (donors with at least 10 junction chondrocytes) were correlated with grade by Spearman’s rank correlation (nominal *p*); the differential expression and GSEA of the state are those of the per-state model above (capture-burden covariate recomputed on the junction chondrocytes). For the snATAC, the barcodes of the junction chondrocytes were matched to the ATAC fragments and the consensus peak-by-cell matrix, pseudobulked per donor (14 donors with at least 10 nuclei) and scored for motif activity as described below. For peak-level analysis, log2 CPM of each of the 138,660 consensus peaks above the detection floor was correlated with grade across the 13 donors of the consensus peak-by-cell matrix; opening and closing sets were defined at nominal *p* < 0.05. Chondrocyte-specific peaks were the 5,000 peaks with the largest difference in mean log2 CPM between the junction chondrocytes and all other bone cells. Motif occurrence per peak was taken from the Cluster-Buster top-1% peak sets; for each motif, enrichment of a peak set was tested by a hypergeometric test against all tested peaks, with Benjamini-Hochberg correction across motifs within each set, and motifs were grouped into families by their directly annotated transcription factors; the best motif per family is shown. Peaks within 250 kb of *FGF1*, *FGF2* and *FN1* were analyzed per peak as for the osteocyte loci: linear slope of log2 CPM on grade, Spearman *p* (nominal), and correlation with the junction-chondrocyte RNA of the gene across the same donors (linked when rho ≥ 0.5 and *p* < 0.05).

### Cell-cell communication

CellChat was run separately on each stage (Normal, early OA, severe OA) over the same 26 cell states, the chondrocyte state restricted to the junction chondrocytes, on library-size normalized, log-transformed counts, with the human ligand-receptor database excluding non-protein signaling. A state required 25 nuclei within a stage to be included. Communication probabilities were computed with the triMean method and population.size = TRUE; communications supported by fewer than 10 cells were filtered out, and pathway-level probabilities were computed from the retained interactions (Supplementary Tables 9 to 11). Because single-nucleus data under-represent cytoplasmic transcripts, secreted ligands are detected less efficiently than in whole-cell data; comparisons between stages are on equal footing, but absolute interaction counts are not comparable with scRNA-seq reports. Circle plots share one edge-weight maximum per pathway across the three stages. The osteocyte’s share of TGF-β signaling at each stage is its summed outgoing communication probability in the TGFb pathway divided by the pathway total over all sending states; receptor shares of the FN1 input onto the osteocyte were computed analogously over the FN1 ligand-receptor pairs. Donors were pooled within a stage, so the communication probabilities carry no donor-level replication; the CellChat results are descriptive and are reported without inferential statistics.

### Trajectory and pseudotime

Slingshot was run on the first 20 principal components of the integrated atlas embedding for eight clusters: the five osteogenic states (osteocyte states merged) together with the CAR cell and the activated and stress fibroblasts, with the CAR cell as start cluster, based on the lineage evidence that osteoblasts arise from LepR+/*CXCL12*+ marrow stromal cells. The analysis was also run without specifying a start cluster and returned the same ordering of the CAR cell upstream of the osteoprogenitor. Pseudotime is the average across lineages (Supplementary Table 8). Marker and receptor genes were fitted along pseudotime as loess curves (span 0.5, degree 1) of per-nucleus log-normalized expression over the 19,063 lineage nuclei, min-max scaled per gene for display. The osteocyte segment of the trajectory was defined as pseudotime above 85 and the osteoblast segment as 60 to 85, from the median pseudotime of each state; segment enrichment is the ratio of mean unscaled fitted expression between the two segments. For the *FN1* axis display, *FN1* was quantified in the chondrocyte pseudobulk and its receptors (*CD44*, *ITGAV*) in the merged-osteocyte pseudobulk as log2 counts per million per donor; the nominal p values shown are those of the per-state continuous-grade model, whose adjusted values are given in the Results, with no statistics re-estimated for the display.

### Single-nucleus ATAC processing

The 16 libraries were generated on the 10x Chromium Single Cell Multiome ATAC + Gene Expression platform, so every nucleus carries a paired transcriptome and accessibility profile. ATAC reads were processed with Cell Ranger ARC, and the per-sample fragment files were used for all analyses. Nuclei were assigned to cell states by transferring the frozen snRNA-seq annotations through the shared nucleus barcode (all 62,993 labeled nuclei matched their fragment-file barcodes); no clustering was performed on the ATAC data. Per-nucleus fragment counts and the fraction of fragments in peaks were computed for quality control; the osteogenic states showed the highest fraction of fragments in peaks in the dataset.

### Peak calling and the consensus peak set

Fragments were pooled per cell state and peaks were called per state with MACS3 (v3.0.1; --nomodel --shift -75 --extsize 150 --keep-dup all -q 0.05 --call-summits), so that peaks of rare states such as the osteocyte are not diluted by the abundant states. A consensus peak set was then built from the state-level summits: each summit was extended to a fixed 500-bp window, candidates were ranked by MACS3 score, and windows were retained greedily so that no two retained peaks overlap, yielding 283,535 consensus peaks. Fragment counts per nucleus over the consensus peaks were aggregated into donor-by-state pseudobulk profiles.

### Transcription-factor motif activity

The consensus peak sequences were scored against the 10,249 motifs of the SCENIC+ v10nr_clust collection with Cluster-Buster, using the create_cisTarget_databases framework, giving a motif-by-peak score matrix. For the chondrocyte, the donor-by-state pseudobulks were rebuilt from the junction chondrocytes (14 donors with at least 10 nuclei) and scored in the same way. For each motif, the top 1% of peaks by score were taken as that motif’s peak set, and motif activity in a donor-by-state pseudobulk was defined as the fraction of its fragments falling in the motif’s peak set, divided by the fraction of fragments in all peaks (depth normalization). Donor-by-state groups with fewer than 10 nuclei were excluded, and a state was shown only if 10 such donors remained, the rule used for the expression analysis. Motifs were assigned to transcription-factor families by their database annotations; for a family score, each member motif’s activity was z-scored across the donors of a state and the member z scores were averaged.

### Statistics of the chromatin figures

Family activity was compared between severe OA and Normal and early OA donors within each state by two-sided Wilcoxon rank-sum test and reported at nominal p (**Figure 5a, b**), matching the convention of the composition test; Supplementary Table 12 gives, alongside, the Benjamini-Hochberg adjusted p within each family across the 25 tested states and the Spearman correlation of the family score with continuous grade with its nominal and adjusted p. Other associations with continuous grade and between donor-level quantities (motif activity, pseudobulk expression) were tested by Spearman correlation at nominal p, and the composition test (**Figure 7a**) is reported at nominal p as before. For **Figure 7e, t**he same Spearman test was run per motif for all 562 factors with an annotated motif and reported for the osteogenic and adipogenic factors shown; this scan is exploratory and its p values are nominal. The marker and receptor curves along pseudotime (Figure 7c and Figure 8d) are loess fits (span 0.5, degree 1) of log-normalized expression against the CAR-start pseudotime, min-max scaled per gene. For Figure 8, osteocyte pseudobulk expression was computed per donor from the nuclei of the merged osteocyte state, as counts per million, over the 16 donors of the per-state model; correlations with junction chondrocyte FN1 used the 14 donors with an eligible junction chondrocyte pseudobulk. The osteocyte motif activity of Figure 8a rests on the same 16 donors. The osteocyte panels (Figure 8b and c) report the per-state grade model of Figure 4 (nominal p) and donor-level Spearman correlations against the osteocyte SMAD2/3 family activity; Figure 8f is the Spearman correlation of the lineage-averaged SMAD2/3 score with chondrocyte FN1.

### Statistics and reproducibility

All statistical tests are described in their respective sections; tests are two-sided unless stated otherwise. All analyses were performed in R; every figure panel is generated by a versioned script from frozen intermediate outputs, and no statistics are re-estimated at figure time.

## Data availability

Raw and processed sequencing data analyzed in this study (bulk RNA-seq and single-nucleus multiome RNA-seq and ATAC-seq) are derived from the donors included in datasets that will be available through the SPARC Portal (RRID:SCR_017041) upon publication at the following DOIs: https://doi.org/10.26275/lyd6-v13s (bulk RNA-seq)^81^ and https://doi.org/10.26275/k5zn-h5y3 (single-nucleus multiome)^82^. Raw sequencing data will be available through dbGaP under accession phs004628. Tissue-processing protocols are available at protocols.io (https://dx.doi.org/10.17504/protocols.io.j8nlk8mrdl5r/v1; https://dx.doi.org/10.17504/protocols.io.6qpvr8m2zlmk/v1). All processed results underlying the figures are provided as Supplementary Tables 1 to 14.

## Software

Histological images were analyzed in QuPath v0.5.1. Bulk RNA-seq reads were aligned with HISAT2; gene-level differential expression was fitted with DESeq2 and gene set enrichment was run with clusterProfiler. Transcript abundances were estimated with StringTie v1.2.3 against GENCODE v44 and isoform usage was analyzed with IsoformSwitchAnalyzeR. Single-nucleus multiome reads were processed with Cell Ranger ARC (10x Genomics, GRCh38-2020-A). The single-nucleus RNA-seq data were normalized, integrated and clustered with Seurat v5; per-state differential expression was fitted with DESeq2 on pseudobulk counts, intercellular communication was inferred with CellChat, and trajectories were fitted with Slingshot. Peaks were called from the single-nucleus ATAC fragments with MACS3 v3.0.1. All statistics and figures were produced in R.

## Acknowledgements

The authors thank Michelle Payad, Zaira Flores, Jessica Nitta, and the team at Lifesharing, San Diego for providing access to the precious gift of tissue donation from the local San Diego and Imperial communities that made the research possible. We thank the Scripps Research Genomics core members Tony Mondala, Stephen Head and Sean Wang for their support with omics data generation.

## Author contributions

M.K.L., Y.U., H.S. and H.A. conceptualized this study. Y.U., H.S. and M.K.L. drafted the manuscript, with contributions from the other authors. Y.U. and H.S. performed data analysis. All authors approved the final version of the manuscript and figures.

## The RE-JOIN Consortium

Armen Akopian, Kyle Allen, Alejandro Almarza, Benjamin Arenkiel, Yangjin Bae, Bruna Balbino de Paula, Anita Bandrowski, Mario Danilo Boada, Jacqueline Boccanfuso, Jyl Boline, Dawen Cai, Dellina Lane Carpio, Robert Caudle, Racel Cela, Yong Chen, Rui Chen, Brian Constantinescu, Ibdanelo Cortez, Yenisel Cruz-Almeida, M. Franklin Dolwick, Chris Donnelly, Zelong Dou, Joshua Emrick, Malin Ernberg, Danielle Freburg-Hoffmeister, Spencer Fullam, Janak Gaire, Akash Gandhi, Benjamin Goolsby, Stacey Greene, Nele Haelterman, Michael Iadarola, Shingo Ishihara, Azeez Ishola, Sudhish Jayachandran, Zixue Jin, Frank Ko, Priya Kulkarni, Zhao Lai, Brendan Lee, Yona Levites, Carolina Leynes, Jun Li, Martin Lotz, Lindsey Macpherson, Tristan Maerz, Camilla Majano, Anne-Marie Malfait, Maryann Martone, Bella Mehta, Richard Miller, Rachel Miller, Michael Newton, Alia Obeidat, Merissa Olmer, Dana Orange, Miguel Otero, Kevin Otto, Folly Patterson, Marlena Pela, Sienna Perry, Theodore Price, Hernan Prieto, Russell Ray, Dongjun Ren, Margarete Ribeiro Dasilva, Alexus Roberts, Elizabeth Ronan, Oscar Ruiz, Shad Smith, Mairobys Socorro, Kaitlin Southern, Joshua Stover, Michael Strinden, Hannah Swahn, Evelyne Tantry, Sue Tappan, Luis Tovias Sanchez, Airam Vivanco-Estela, Joost Wagenaar, Lai Wang, Kim Worley, Joshua Wythe, Jiansen Yan, and Julia Younis.

## Supplementary Information

### Supplementary Figures

**Supplementary Figure 1.**
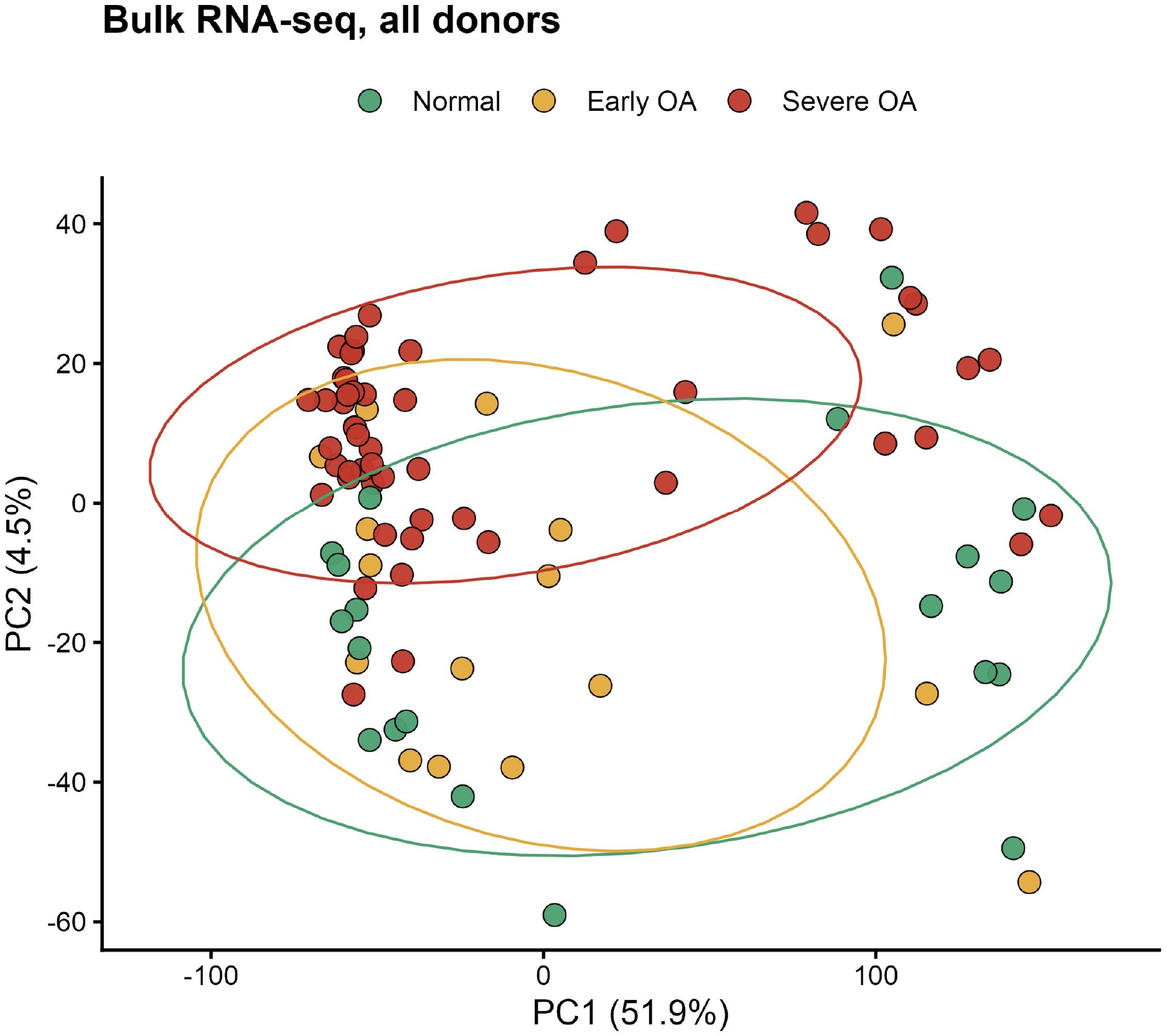
Principal component analysis of the bulk RNA-seq cohort. The 89 donors on the first two principal components of the 2,000 most variable genes, colored by stage; ellipses are 68% normal contours. Details in Methods.

**Supplementary Figure 2.**
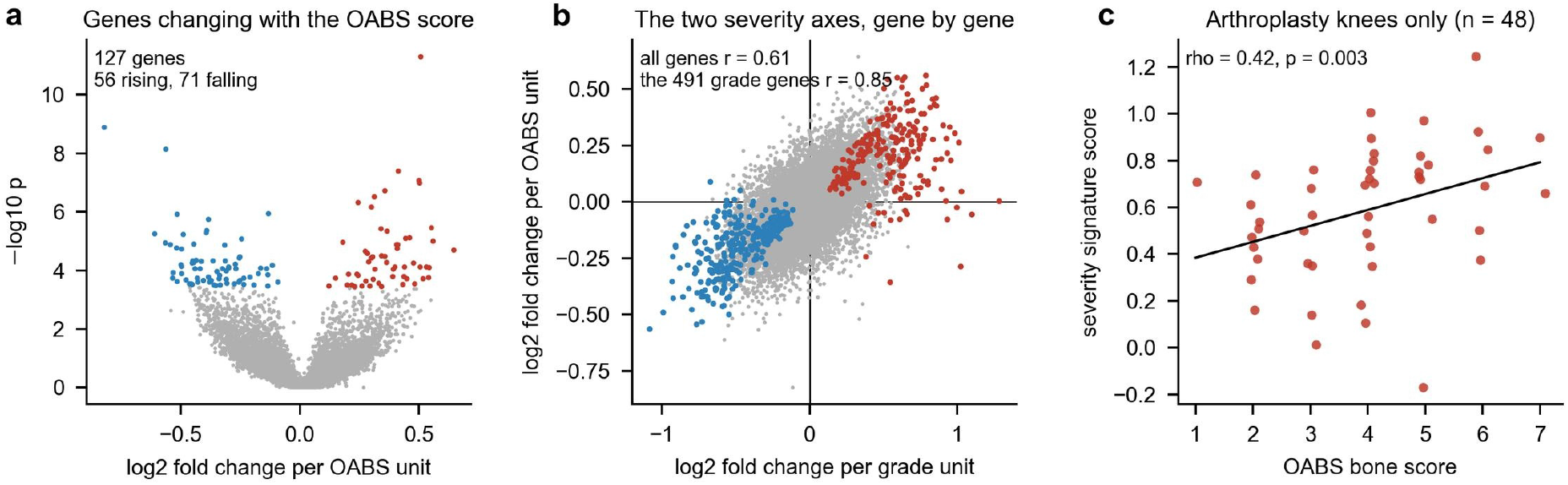
The OABS bone score as a second severity axis. (a) Genes changing with the OABS score of the medial weight-bearing subchondral bone across the 85 knees that carry one, tested under the model of the macroscopic grade; red rising, blue falling, Benjamini-Hochberg adjusted p below 0.05 over the 17,366 tested genes. (b) Log2 fold change per grade unit against log2 fold change per OABS unit over the same genes; the 491 genes that change with grade are colored by the direction of their grade effect. (c) The severity signature of the 491 grade genes, the mean z score of the 224 rising genes minus the mean of the 267 falling genes, against the OABS score within the 48 arthroplasty knees, where the macroscopic grade is uniform at 4; points are jittered horizontally, the line is a least-squares fit, and Spearman rho with nominal p is given. Details in Methods.

**Supplementary Figures 3 to 6.**
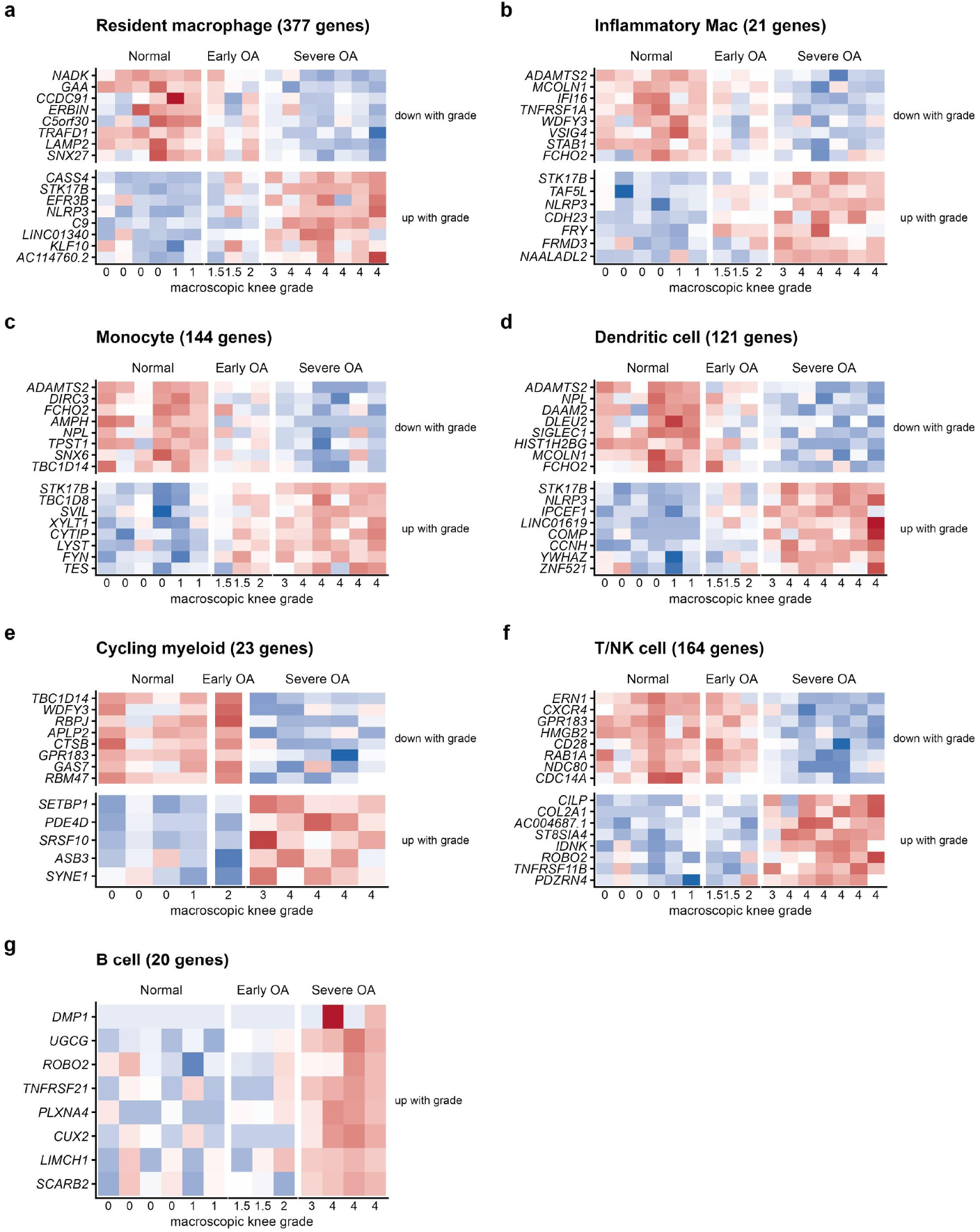
Genes changing linearly with grade in the remaining cell states. Per-state pseudobulk heatmaps as in Figure 4f to i, for the myeloid and lymphoid states (Supplementary Fig. 3), the vascular and perivascular states (Supplementary Fig. 4), the marrow stromal states (Supplementary Fig. 5) and the WIF1+ osteoblast and the mature osteoblast (Supplementary Fig. 6). States shown in Figure 4 do not reappear here. The plasma cell and the stress fibroblast have three or fewer significant genes, and the osteoclast and the early osteocyte fall below the ten-donor threshold of the per-state model; these states are reported in the tables only. Details in Methods.

**Supplementary Figure 4.**
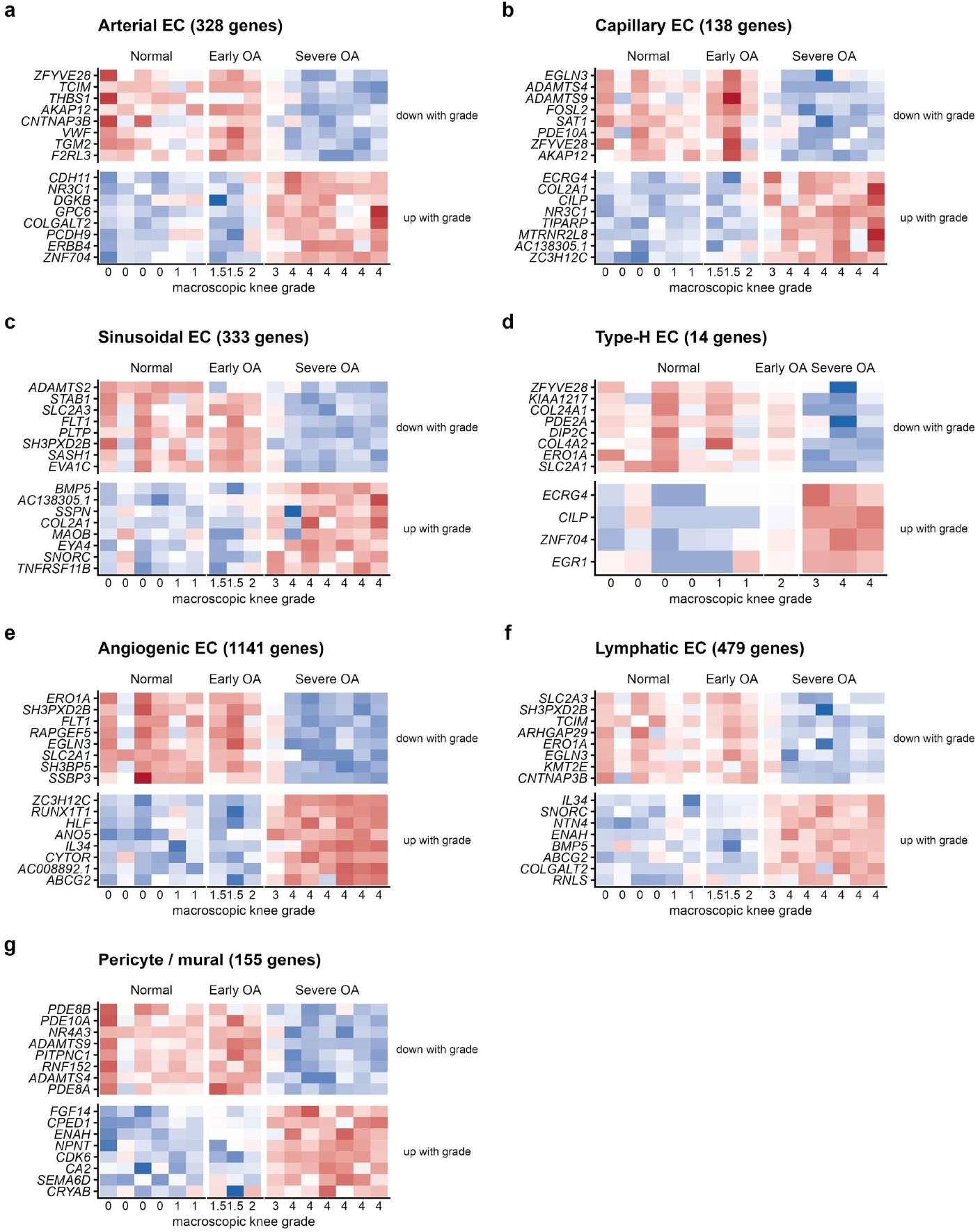
Genes changing linearly with grade in the vascular and perivascular states (legend with Supplementary Figure 3).

**Supplementary Figure 5.**
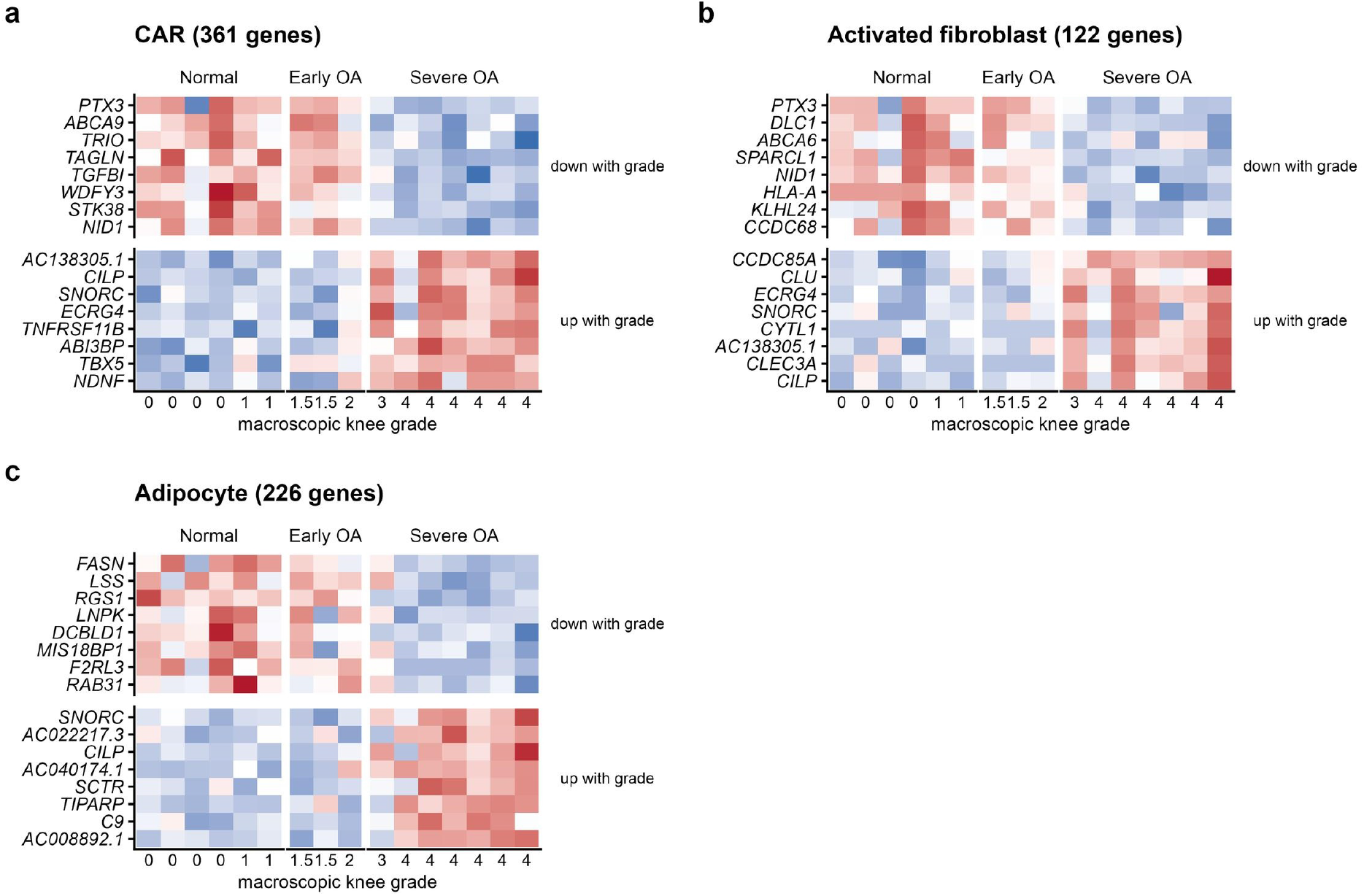
Genes changing linearly with grade in the marrow stromal states (legend with Supplementary Figure 3).

**Supplementary Figure 6.**
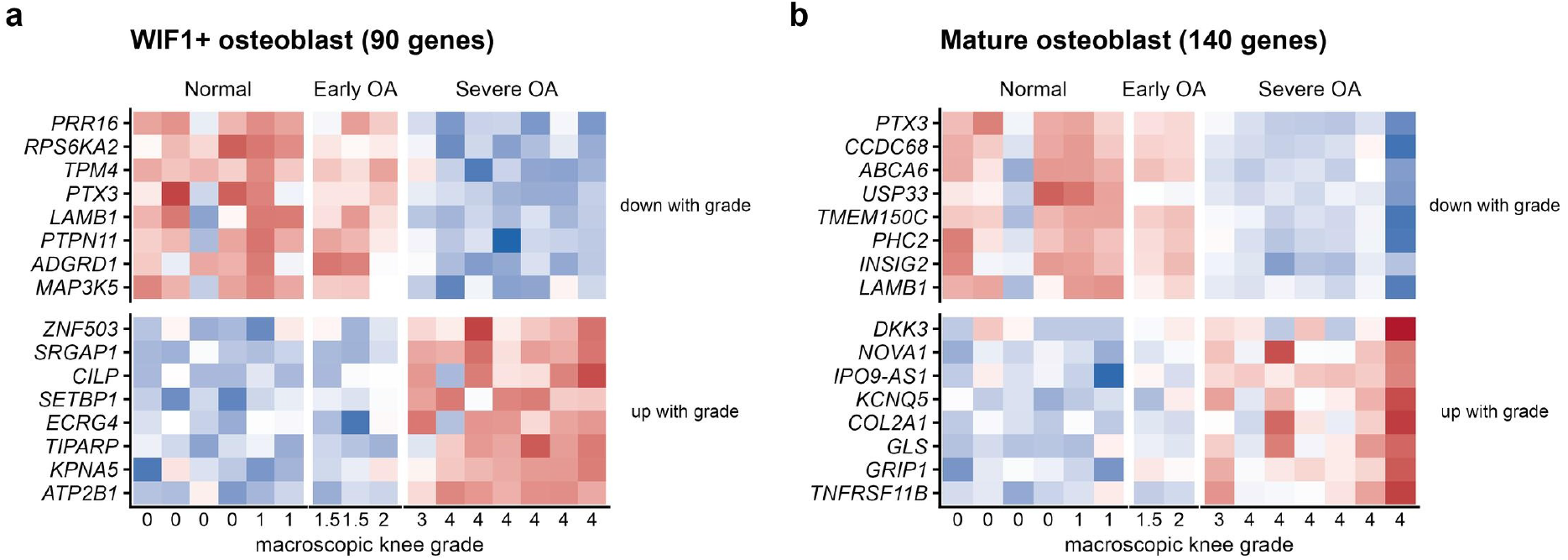
Genes changing linearly with grade in the chondrogenic and osteogenic states (legend with Supplementary Figure 3).

**Supplementary Figure 7.**
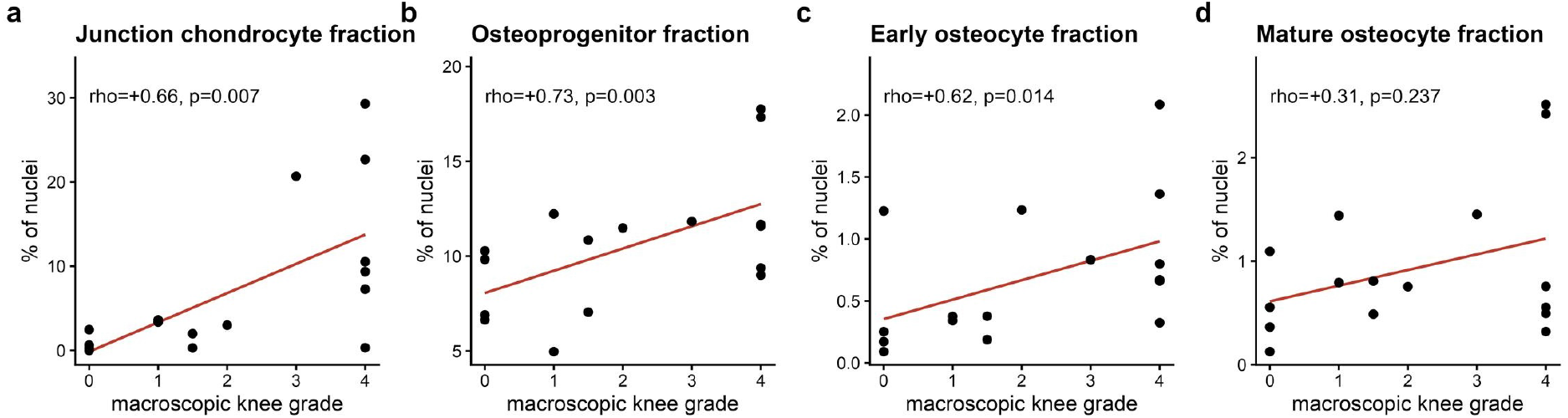
Junction chondrocyte and osteoprogenitor fractions against grade. Share of nuclei per donor for (a) the junction chondrocyte, (b) the osteoprogenitor, (c) the early osteocyte and (d) the mature osteocyte, displayed as the merged osteocyte in Figure 7b; lines are least-squares fits and the annotations give the abundance test of Figure 7a (for c and d, the same test run on the 27 labels with the two osteocyte states separate).

### Supplementary Table legends

**Supplementary Table 1. Donor information, bulk cohort.** The 89 bulk RNA-seq donors with macroscopic knee grade, the harmonized stage assignment, age, sex, BMI, post-mortem interval and sequencing quality metrics.

**Supplementary Table 2. Donor information, single-nucleus cohort.** The 16 single-nucleus donors with grade, stage, age, sex, BMI, post-mortem interval and nuclei counts.

**Supplementary Table 3. Bulk RNA-seq differential expression against continuous macroscopic knee grade.** All 17,366 tested genes from the DESeq2 model of the run of record; sig marks Benjamini-Hochberg adjusted *p* < 0.05 (491 genes).

**Supplementary Table 4. Bulk RNA-seq gene set enrichment against grade.** All significant GO biological process terms (BH-adjusted *p* < 0.05; 265 terms) from GSEA on the continuous-grade ranking, with their leading-edge genes.

**Supplementary Table 5. Isoform usage in bulk RNA-seq.** Isoform fraction against continuous macroscopic knee grade for the 90,378 tested isoforms of 14,100 genes: slope in percentage points per grade unit from the model with batch and the detected-genes covariate, the partial Spearman p and its BH-adjusted value (first sheet); the enrichment of isoform classes among rising and falling isoforms shown in Fig. 2e (second sheet); and the isoforms of Fig. 2f (third sheet).

**Supplementary Table 6. Single-nucleus differential expression per cell state.** All 5,654 significant gene-by-cluster associations (adjusted p < 0.05; each gene counted once per cluster in which it changed) from the per-state pseudobulk model with the capture-burden covariate, across the 25 cell states that clear the donor threshold, with the donor count of each state.

**Supplementary Table 7. GSEA of the osteogenic states against grade.** All significant GO biological process terms (BH-adjusted *p* < 0.05) from GSEA on the continuous-grade ranking of the osteocyte, the osteoprogenitor, the *WIF1*+ osteoblast and the stromal osteoblast (433 terms).

**Supplementary Table 8. Trajectory analysis.** Per-nucleus pseudotime for the 19,063 nuclei of the eight stromal-osteogenic states (CAR-start Slingshot), the lineages returned by the CAR-start and unsupervised runs, and the loess fits of the marker cascade and of receptor genes along pseudotime.

**Supplementary Tables 9 to 11. CellChat ligand-receptor interactions.** All inferred interactions with their communication probabilities in Normal (Supplementary Table 9), early OA (Supplementary Table 10) and severe OA (Supplementary Table 11), with pathway assignment and database evidence.

**Supplementary Table 12. Transcription-factor family motif activity in single-nucleus ATAC.** Transcription-factor family motif activity for every family and cell state of Figure 5a: the number of donors, the severe OA minus Normal and early OA difference of family z scores with its nominal Wilcoxon p (the stars of Figure 5a, b) and BH-adjusted p (within each family across the 25 tested states), and the Spearman correlation of the donor-level family score with continuous macroscopic knee grade with its nominal and BH-adjusted p. Donor-by-state groups with at least 10 nuclei and states with at least 10 such donors. The second sheet lists the pseudobulk groups.

**Supplementary Table 13. Composition against grade.** Spearman correlation of each state’s centered log-ratio abundance with continuous grade, donor-label permutation *p* and BH-adjusted *p* across the 26 states (first sheet), the same test on the 27 labels with the two osteocyte states separate, and the pre-specified lineage tests.

**Supplementary Table 14. Junction chondrocytes.** The quality filter of the chondrocyte label by stage and donor; markers of the nine integrated subclusters; subcluster shares per donor and their correlation with grade; pseudobulk log2 CPM per donor and its correlation with grade; differential expression against grade (the model of record with the capture-burden covariate, and the model without it); GSEA; specificity against the other 25 states; snATAC motif activity against grade; motif enrichment in the chondrocyte-specific, opening and closing peak sets (all motifs, and the best motif per family); and the peaks within 250 kb of *FGF1*, *FGF2* and *FN1*.

